# Top-down control of flight by a non-canonical cortico-amygdala pathway

**DOI:** 10.1101/2022.05.19.492688

**Authors:** Chandrashekhar D. Borkar, Claire E. Stelly, Xin Fu, Maria Dorofeikova, Quan-Son Eric Le, Rithvik Vutukuri, Catherine Vo, Alex Walker, Samhita Basavanhalli, Anh Duong, Erin Bean, Alexis Resendez, Jones G. Parker, Jeffrey G. Tasker, Jonathan P. Fadok

## Abstract

Survival requires the selection of appropriate behaviour in response to threats, and dysregulated defensive reactions are associated with psychiatric illnesses such as posttraumatic stress and panic disorder.^1^ Threat-induced behaviours, including freezing and flight, are controlled by neuronal circuits in the central amygdala (CeA)^2^; however, the source of neuronal excitation to the CeA that contributes to high-intensity defensive responses is unknown. Here we used a combination of neuroanatomical mapping, *in-vivo* calcium imaging, functional manipulations, and electrophysiology to characterize a previously unknown projection from the dorsal peduncular (DP) prefrontal cortex to the CeA. DP-to-CeA neurons are glutamatergic and specifically target the medial CeA, the main amygdalar output nucleus. Using a behavioural paradigm that elicits both freezing and flight, we found that CeA-projecting DP neurons are activated by high-intensity threats in a context-dependent manner. Functional manipulations revealed that the DP-to-CeA pathway is necessary and sufficient for both avoidance behaviour and flight. Furthermore, we found that DP projections target distinct medial CeA neuronal populations projecting to midbrain flight centres. These results elucidate a non-canonical top-down pathway regulating defensive responses.

---

In the face of threat, organisms display a continuum of defensive behaviours and flexibly shift between defensive strategies.^3,4^ Selecting the appropriate action for survival depends on the proximity and intensity of the threat and the context of the encounter.^5,6^ Moreover, exaggerated responses to perceived threats have been associated with anxiety, post-traumatic stress and panic disorders, and phobias.^1,7,8,9^ Abnormal patterns of activation in the prefrontal cortex (PFC) are associated with these mental illnesses, and bidirectional projections between the medial PFC (mPFC) and basolateral amygdala are part of a canonical pathway that has been extensively studied in the acquisition and expression of learned fear measured with freezing.^10–14^ Previous studies have shown that animals exhibit behavioural scaling to favour flight over freezing in response to a high-intensity threat, and conditional freeze-to-flight shifting is regulated by distinct and mutually inhibitory local circuit motifs in the CeA.^2,15,16^ Whether the cortex exerts top-down control of these behavioural shifts is not known. Interestingly, the mPFC also projects to the CeA,^17^ raising the possibility that a direct pathway from mPFC to CeA could influence defensive action selection. However, this pathway has never been defined neuroanatomically, and its role in regulating defensive behaviour is unknown.

## Direct mPFC projections to medial CeA

To determine which subdivision of the mPFC projects to the CeA, we injected fluorescent, retrogradely trafficked latex microspheres into the CeA (**Fig. 1A, B**). We observed sparse projections from both the prelimbic (PL) and infralimbic (IL) cortices. However, a significantly greater number of CeA projectors were located in the dorsal peduncular nucleus (DP; **Fig. 1C, D; Fig. S1A**), a region of the mPFC recently linked to physiological and behavioural responses to stress.^18^

**Figure 1.**
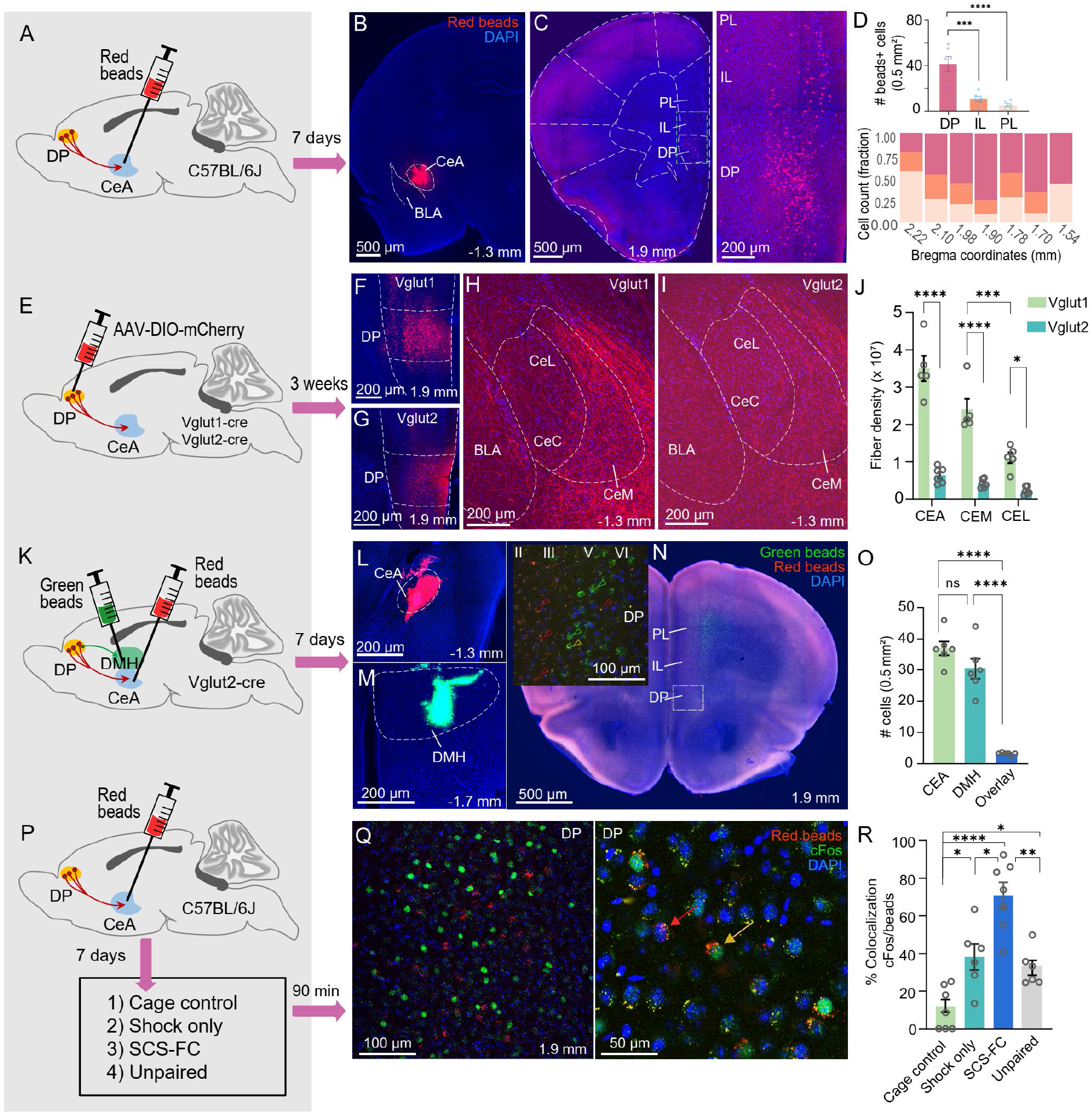
Neuroanatomical characterization of the DP-to-CeA pathway. **A**, Schematic diagram showing retrograde tracing strategy. **B**, Histology image showing injection site of red beads in the CeA. **C**, Histology images of red retro-beads+ neurons in the DP (at 5x *(left)* and 10x *(right)* magnification). **D**, *Top,* A significantly higher proportion of mPFC CeA-projecting neurons are in the DP, compared to the IL or PL (One-way ANOVA, F_(2,15)_ = 21.86, *p* < 0.0001; Bonferroni’s post-hoc test, \*\*\**p*<0.001, **** *p*<0.0001; N = 6). Data represented as means ± s.e.m. *Bottom*, Distribution of CeA-projecting neurons along the mPFC antero-posterior axis (N = 6; 4-6 sections/mouse). DP, dorsal peduncular cortex; IL, infralimbic cortex; PL, prelimbic cortex. **E**, Strategy used to trace glutamatergic terminals in the CeA. **F**, Representative image of DP targeting in Vglut1-Cre mice. **G**, Representative image of DP targeting in Vglut2-Cre mice. **H**, Representative expression of mCherry+ terminals in the CeA of a VGlut1-Cre mouse. **I**, Representative expression of mCherry+ terminals in the CeA of a VGlut2-Cre mouse. **J**, Integrated density of mCherry+ fibres from Vglut1+ (N = 5, 3-4 sections/mouse) and Vglut2+ (N = 6, 2-3 sections/mouse) neurons (unpaired Student’s t-test for total CeA, Vglut1 vs Vglut2, t = 8.89, df = 9, *p* < 0.0001. Two-way ANOVA, CeM vs CeL, strain X region, F_(1,18)_ = 12.6, *p* < 0.01; Bonferroni’s post-hoc test, \**p* < 0.05, \*\*\*\**p* < 0.0001. CeL includes lateral and capsular subregions. Data represented as means ± s.e.m. **K**, Dual-target retrograde tracing strategy from CeA and dorsomedial hypothalamus (DMH). **L**, Representative targeting of CeA. **M**, Representative targeting of DMH. **N**, Deposition of green and red beads in the mPFC. *Inset,* distribution of CeA and DMH projectors in different layers of DP. Red, green, and yellow arrowheads indicate red beads, green beads, and overlay, respectively. **O**, Number of DP neurons projecting to CeA, DMH, or both (One-way ANOVA, F_(2,15)_ = 62.06, *p* = 0.0001; Bonferroni’s post-hoc test, \*\*\*\**p* < 0.0001; N = 6, 3 sections/mouse). Data represented as means ± s.e.m. **P**, Strategy for neuronal activation analysis. Seven days after injection, animals were split into four groups: home cage control, shock only control, unpaired control, and serial compound stimulus fear conditioning (SCS-FC). **Q**, Representative images showing bead+ and/or cFos+ cells in the DP. *Left*, 10x and *Right*, 20x magnifications. Red arrow, bead+; yellow arrow, bead and cFos+. **R**, Quantification of cFos expression in bead+ cells (N = 6-7 mice; 3-4 sections/mouse; One-way ANOVA, F_(3,22)_ = 14.47, *p* < 0.0001; Bonferroni’s post-hoc test, \**p* < 0.05, \*\**p* < 0.001, \*\*\*\**p* < 0.0001). Data represented as means ± s.e.m.

The mPFC regulates cognition, motivation, and emotion via distinct glutamatergic projections to cortical and subcortical regions.^19^ The vesicular glutamate transporter (Vglut) 1 is used to label most excitatory cortical neurons, yet the DP is enriched in both the Vglut1 and Vglut2 isoforms, which define different populations of projection neurons.^20^ To characterize the DP-to-CeA pathway more precisely, we injected a Cre-dependent mCherry vector into the DP of Vglut1- or Vglut2-Cre mice and quantified fibre density in CeA subnuclei (**Fig. 1E–G**). We found that, while the mCherry+ fibre density within the CeA was significantly higher in Vglut1 mice (**Fig. 1H–J**), notably, in both Vglut1 and Vglut2 mice, mCherry+ fibres were most abundant in the medial subdivision of the CeA (CeM), as compared to the lateral (CeL) or capsular (CeC) subdivisions (**Fig. 1J**). Therefore, Vglut1- and 2-expressing DP neurons are anatomically positioned to influence the function of the primary amygdalar output centre controlling the expression of adaptive behaviour.

The DP also projects to the dorsomedial hypothalamus (DMH), a pathway known to regulate sympathetic responses to psychosocial stress.^18^ To determine whether the DP-to-CeA and DP-to-DMH pathways are discrete, we injected red or green-fluorescent retro-beads into the CeA and DMH (**Fig. 1K–M**). Both red and green retro-beads were localized in DP cells (**Fig. 1N**), but the DMH-projecting DP neurons were localized in layers V and VI (*90% vs 8.7%* in layer II/III), whereas CeA-projecting cells were found mostly in layers II/III (**Fig. 1N, *inset;* Fig. S1B**, *87% vs 10.5%* in layers V and VI). Only a small number of DP cells (∼7%) were co-labelled, suggesting that the DP-to-CeA and DP-to-DMH pathways are anatomically distinct (**Fig. 1O; Fig. S1B**).

Given the role of the CeA in defensive response selection and the role of the DP in stress responding,^2,18^ we hypothesized that threats would activate CeA-projecting DP neurons. To test this, we injected C57BL/6J mice with retro-beads in the CeA and divided animals into four groups: home cage control, footshock-only, unpaired, and fear conditioning (**Fig. 1P**). The fear conditioning (FC) group was subjected to a protocol we developed that elicits freezing and flight responses to separate components of a serial compound stimulus (SCS-FC; **Fig. S1D–E**).^2^ Compared to home cage control, shock-only, and unpaired groups, SCS-FC mice displayed significantly greater numbers of retro-bead+ cells in the DP that were co-labelled with cFos (**Fig. 1Q, R**). There were no significant differences in the total number of retro-bead+ cells across the groups, indicating a consistent retrograde labelling efficiency across the experimental groups (**Fig. S1C**).

## DP-to-CeA neurons are activated by flight

The significantly elevated expression of cFos in the SCS-FC group suggests that DP-to-CeA neurons are recruited most under conditions that elicit robust cue-induced defensive behaviour. Therefore, we set out to determine the activation patterns of DP-to-CeA projecting neurons during our SCS-FC protocol, which induces escalating intensities of defensive behaviours (**Fig. S1D, E**). We used an intersectional approach to selectively express GCaMP6f in DP-to-CeA projection neurons and imaged *in-vivo* calcium activity in freely moving mice using custom-built open-source miniscopes (**Fig. 2A–C)**. We observed low levels of freezing and no flight before conditioning (**Fig. S2A–D**), with similar levels of neuronal activation to the tone and white noise (WN) components of the SCS (**Fig. S2E–J**). Moreover, there was no correlation between calcium activity and movement speed before conditioning (**Fig. S2G**). Following conditioning, mice exhibited elevated levels of freezing to the tone, and flight to the WN in the conditioning context (hereafter called the high-threat context; **Fig. 2D–E; Fig. S3A–B**).^2,21^ By contrast, mice exhibited predominantly freezing when exposed to the same conditioned stimuli in a neutral context (hereafter called the low-threat context; **Fig. 2M–N; Fig. S3I–J**). Interestingly, as conditioning progressed, the mean activity of CeA-projecting DP neurons increased concurrently, which correlated significantly with speed in the high-but not low-threat context (**Fig. S3C, K**).

**Figure 2:**
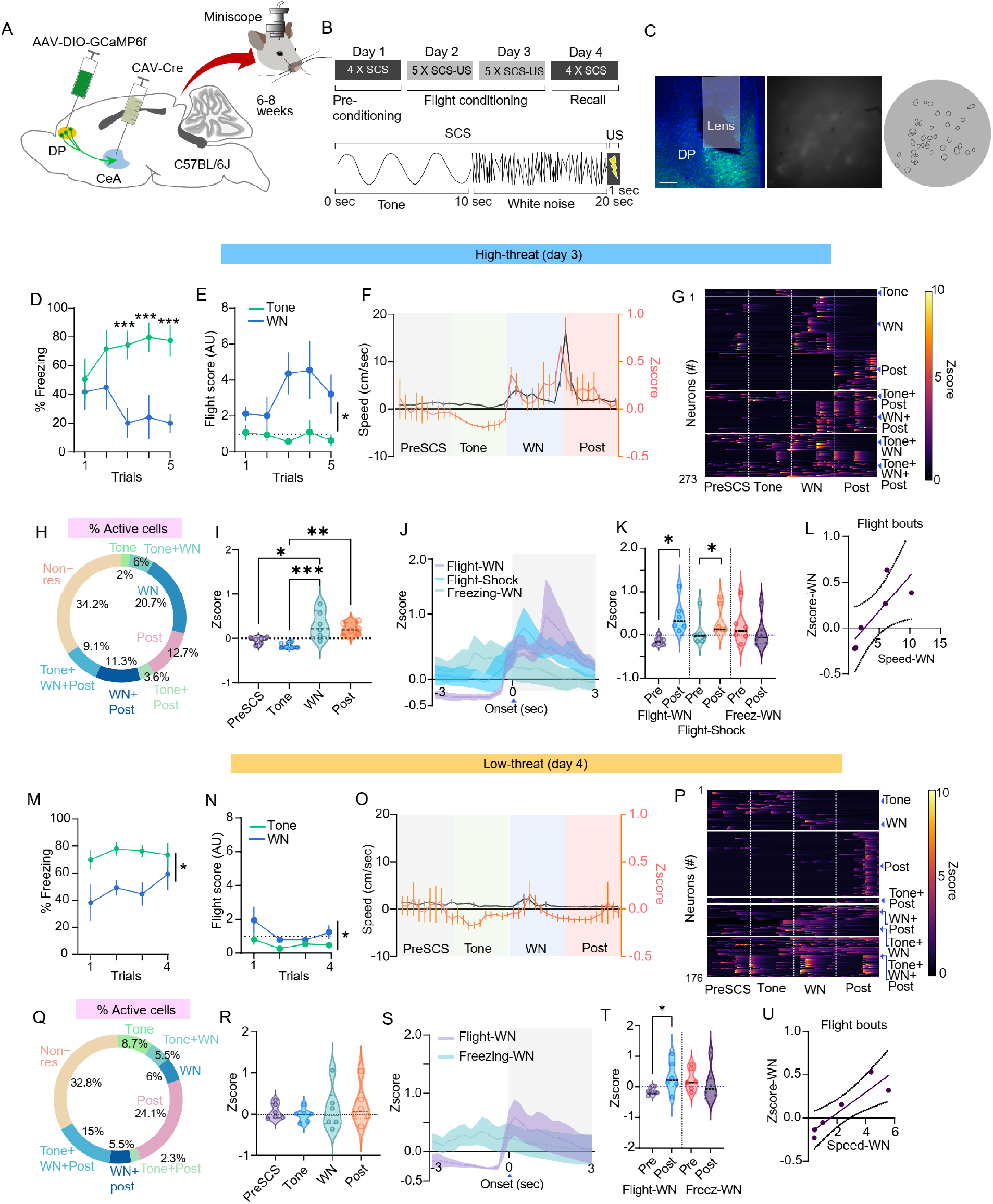
DP-to-CeA projecting cells are activated by high fear states. **A**, Schematic of the intersectional strategy used to record DP-to-CeA projector activity (N = 6 mice). **B**, Implanted mice were subjected to a fear conditioning paradigm designed to elicit conditioned freezing and flight. **C**, *Left*, Representative GCaMP6f expression with lens placement in the DP (scale bar = 500 µm). *Right*, miniscope field-of-view (raw and post cell extraction). **D,** Trial-wise freezing in the high-threat context. Two-way ANOVA, Trial x Stimuli, F_(4, 20)_ = 4.354, *p* = 0.01, followed by Bonferroni’s post-hoc test. \*\**\*p* < 0.001. Data shown as means ± s.e.m. **E,** Trial-wise flight in the high-threat context. Two-way ANOVA, Trial x Stimuli, F_(4, 20)_ = 2.08, *p* = 0.12, Stimuli, F_(1, 5)_ = 13.49, *p* = 0.014, Trial, F_(4, 20)_ = 0.933, *p* = 0.46. Data shown as means ± s.e.m. **F**, Z-scored neuronal activity and speed on the last trial in the high-threat context (n = 273 neurons). Data shown as means ± s.e.m. **G**, Heatmap of neuronal activity during the last trial in the high-threat context (273 neurons from 6 mice). **H,** Percentages of neurons activated (neurons with peaks ≥3 s.d. above baseline) during different cue periods in the high-threat context. **I**, Z-score of preSCS, tone and WN and post-shock (post) of the DP-to-CeA population in the high-threat context (from all trials, Ordinary one-way ANOVA, F_(3,20)_ = 9.33, *p* < 0.001, followed by Bonferroni’s post-hoc test, *P < 0.05, **P < 0.005, ***P < 0.001). Violin plots indicate median, interquartile range, and the distribution of individual data points. **J**, Neuronal activity aligned at the onset of WN-induced freezing and flight, and shock-induced flight in the high-threat context. Data shown as means ± s.e.m. **K**, Neuronal population activity 3 s before and after the onset of freezing and flight responses (Paired t-test flight-WN, t=3.28, df=5, *p<0.05; Mann-Whitney test, flight-shock, *p = 0.0411; Paired t-test freezing-WN, t=1.778, df=5, p=0.13). Violin plots indicate median, interquartile range, and the distribution of individual data points. **L**, Spearman correlation of neuronal activity and speed aligned to WN-induced flight bouts (r = 0.94, 95% CI = 0.003864 to 0.1503, *p < 0.05, each point represents one sec). **M,** Trial-wise freezing in the low-threat context. Two-way ANOVA, Trial x Stimuli, F_(3, 15)_ = 0.5806, *p* = 0.63, Stimuli, F_(1, 5)_ = 11.73, *p* = 0.018, Trial, F_(3, 15)_ = 0.83, *p* = 0.49. Data shown as means ± s.e.m. **N,** Trial-wise flight in the low-threat context. Two-way ANOVA, Trial x Stimuli, F_(3, 15)_ = 1.58, *p* = 0.23, Stimuli, F_(1, 5)_ = 8.12, *p* = 0.035, Trial, F_(3, 15)_ = 1.42, *p* = 0.27. Data shown as means ± s.e.m. **O**, Z-scored neuronal population activity and speed on the last trial in the low-threat context. Data shown as means ± s.e.m. **P**, Heatmap of Z-scores from active neurons during the last trial in the low-threat context (176 neurons from 6 mice). **Q,** Percentages of neurons activated (neurons with peaks ≥3 s.d. above baseline) during different cue periods in the low-threat context. **R,** Z-score of preSCS, tone and WN and post-cue (post) of the entire DP-to-CeA population in the low-threat context (from all trials, Ordinary one-way ANOVA, effect of stimuli F_(3,20)_ = 0.39, *p* = 0.75). Violin plots indicate median, interquartile range, and the distribution of individual data points. **S**, Neuronal activity aligned to the onset of WN-induced freezing and flight in the low-threat context. Data shown as means ± s.e.m. **T**, Neuronal activity aligned to the onset of freezing and flight responses (Paired t-test flight-WN, t=2.58, df=5, *p<0.05; flight-post, t=0.8493, df=5, p = 0.43; freezing-WN, t=0.8493, df=5, p=0.43). Violin plots indicate median, interquartile range, and the distribution of individual data points. **U**, Spearman correlation of neuronal activity and speed aligned to WN-induced flight bouts (r = 0.94, 95% CI = 0.04251 to 0.2010, *p < 0.05. Each point represents one sec).

In the high-threat context, the CeA-projecting DP neuronal population was most active during WN and shock (**Fig. 2F–I**), with many individual DP-to-CeA neurons active specifically during sensory epochs eliciting flight behaviour (WN and post-shock periods, 45% of recorded neurons; **Fig. 2G– H; Fig. S3G–H**). Only 2% of cells encoded the tone (**Fig. 2H**) and average neuronal activity was significantly higher during WN and post-shock compared to the tone (**Fig. 2I**). Interestingly, WN-induced activation was diminished in the low-threat context (**Fig. 2O**), neuronal responses were more evenly distributed across sensory epochs (**Fig. 2P–Q**), and no significant difference was observed between neuronal responses to tone and WN (**Fig. 2R; Fig. S3O–P**). Together, these data suggest that DP-to-CeA neuronal activity is context-dependent and is more associated with sensory stimuli that induce higher-intensity threat behaviour.

To determine if DP-to-CEA projector activity is associated with defensive responses, we analysed neuronal activity around the onset of freezing and flight bouts in both contexts. In the high-threat context, neuronal activity was significantly increased at the onset of flight yet was unchanged in response to freezing (**Fig. 2J–K)**. Neuronal activity was also positively correlated with speed, during flight bouts (**Fig. 2L; Fig. S3D**), but not with locomotion surrounding freezing bouts (**Fig. S3E–F**). Flight occasionally occurs in the low-threat context; therefore, we also analysed neuronal responses to flight and freezing onset under these conditions. Just as in the high-threat context, there was a significant increase in activity following flight onset, and a significant correlation between neuronal activation and speed during flight, yet no increase in activity was noted for freezing (**Fig. 2S–U; Fig. S3L–N**).

## DP-to-CeA neurons mediate avoidance

The prevalence of DP-to-CeA neurons active in the high-threat context suggests a functional role for this pathway in negative-valence related behaviours. We first tested this idea using chemogenetic manipulations of DP-to-CeA neurons in standard avoidance paradigms used to assay anxiety-like states: the open field test (OFT) and the elevated plus maze (EPM). We used an intersectional targeting strategy to express either inhibitory (hM4Di-mCherry) or excitatory (hM3Dq-mCherry) DREADDs in the DP-to-CeA pathway, with an mCherry group serving as a CNO control (**Fig. 3A–C**). DREADD-mediated inhibition of the DP-to-CeA pathway increased the time spent and number of entries in the centre zone of the OFT (**Fig. 3D, F, and G**), as well as increasing the number of entries into the open arm of the EPM (**Fig. 3E, I, and J**) without affecting the total distance travelled (**Fig. 3H**). By contrast, chemogenetic stimulation of this pathway did not induce avoidance behaviour in either assay. These findings suggest that the DP-to-CeA pathway is necessary but not sufficient for innate avoidance behaviour.

**Figure 3:**
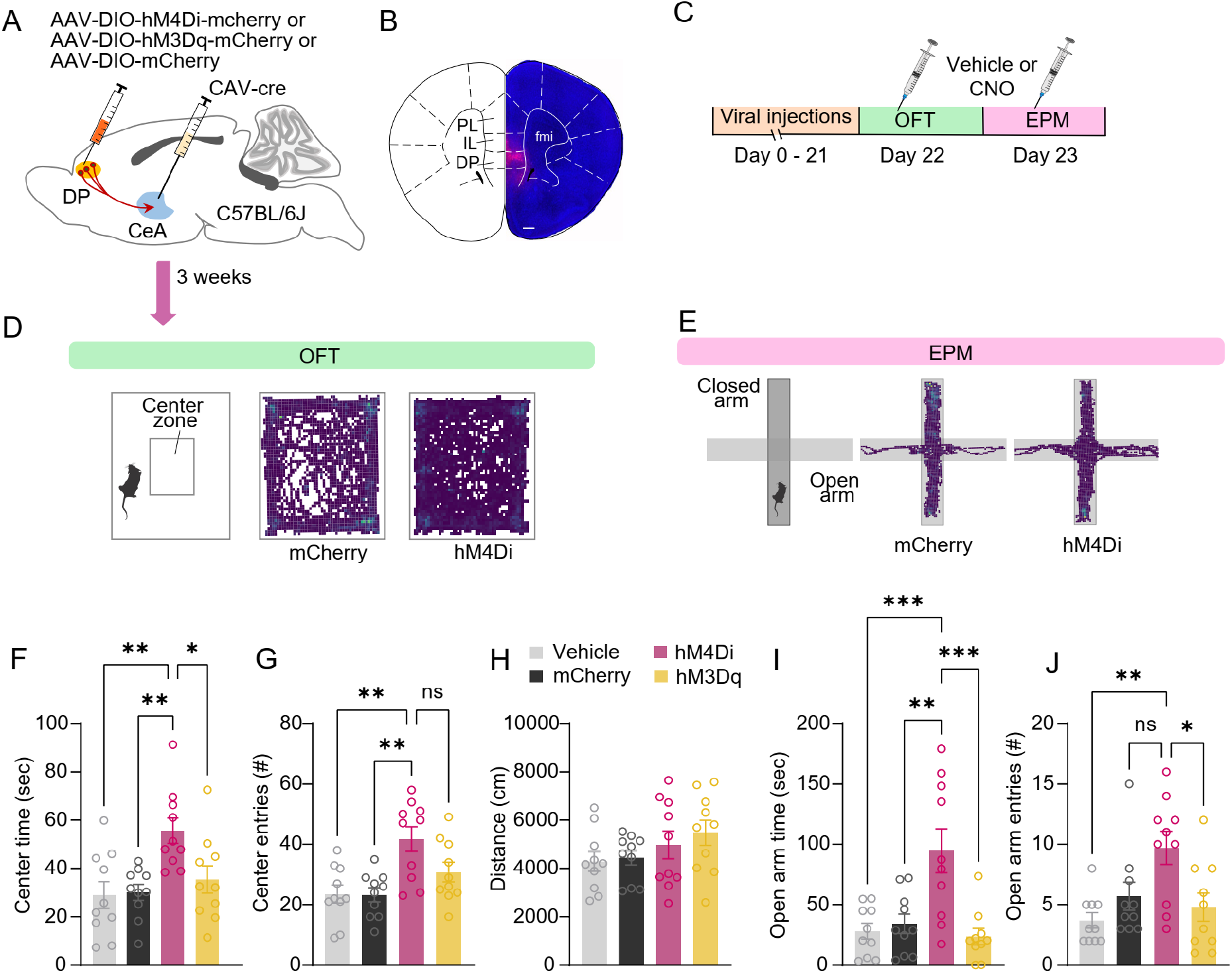
Chemogenetic inhibition of DP-to-CeA pathway reduces avoidance. **A**, Intersectional strategy used for chemogenetic manipulation of the DP-to-CeA pathway. **B**, Representative mCherry expression in the DP (scalebar = 500 µm). **C**, Mice from control (hM4Di-mCherry + vehicle, N = 10; mCherry + CNO, N = 10) and DREADD (hM4Di-mCherry, N = 10; hM3Dq-mCherry, N = 10) groups were subjected to the OFT and EPM 30 min after CNO (5 mg/kg) or vehicle injection. **D**, Representative activity plots of mCherry + CNO and hM4Di + CNO mice in the OFT. **E**, Representative activity plots of mCherry + CNO and hM4Di + CNO mice in the EPM. **F**, Inhibition of DP-to-CeA pathway significantly increased the time spent in the centre zone (One-way ANOVA, F_(3,36)_ = 6.06 *p* = 0.01). Bonferroni’s post-hoc test. \**p* < 0.05, \*\**p* < 0.01. **G**, Inhibition of DP-to-CeA pathway significantly increased the number of entries in the centre zone (One-way ANOVA, F_(3,36)_ = 7.158, *p = 0.0007*). Bonferroni’s post-hoc test. \*\**p* < 0.01. **H**, DREADD manipulations did not alter distance travelled in the OFT (One-way ANOVA, F_(3,36)_ = 1.36, *p* = 0.270). **I**, inhibition of the DP-to-CeA pathway significantly increased open-arm time in the EPM (One-way ANOVA, F_(3,36)_ = 9.22, *p* = 0.0001). Bonferroni’s post-hoc test. \**p* < 0.05; \*\**p* < 0.01; \*\*\**p* < 0.001. **J**, inhibition of the DP-to-CeA pathway significantly increased open-arm entries in the EPM (One-way ANOVA, F_(3,36)_ = 5.47, *p* = 0.003). Bonferroni’s post-hoc test. \**p* < 0.05; \*\**p* < 0.01; \*\*\**p* < 0.001. All values represented as mean ± s.e.m.

## DP-to-CeA neurons mediate flight

To study the necessity of the DP-to-CeA pathway in other forms of defensive behaviour, we performed loss-of-function studies using optogenetics in the OFT and SCS-FC (**Fig. 4 A–F; Fig. S4**). We injected a Cre-dependent vector carrying eNpHR or EYFP control into the DP, and a retrograde CAV2-Cre vector into the CeA of C57BL/6J mice (**Fig. 4A; Fig. S4A**). Inhibition of DP-to-CeA terminals significantly increased the time spent in, and number of entries into, the centre zone of the OFT, but did not affect locomotor activity, consistent with chemogenetic inhibition (**Fig. S4C–E**). We then used a within-subjects experimental design in the SCS-FC paradigm, with optogenetic stimulation during half of the trials (**Fig. S4B**). In the high-threat context, inhibition of DP-to-CeA terminals significantly reduced WN-induced flight (**Fig. 4E; Fig. S4G, H**). Interestingly, inhibiting these terminals reduced freezing to the tone, yet also elevated freezing during the WN period, demonstrating cue-specific reduction in defensive response intensity but not a complete inhibition of fear (**Fig. 4D; Fig. S4F**).^22,23^. Optogenetic inhibition had no effect on defensive behaviour in the low-threat context (**Fig. 4F; Fig. S4I–K)**, further supporting the context-dependent function of the DP-to-CeA pathway.

**Figure 4.**
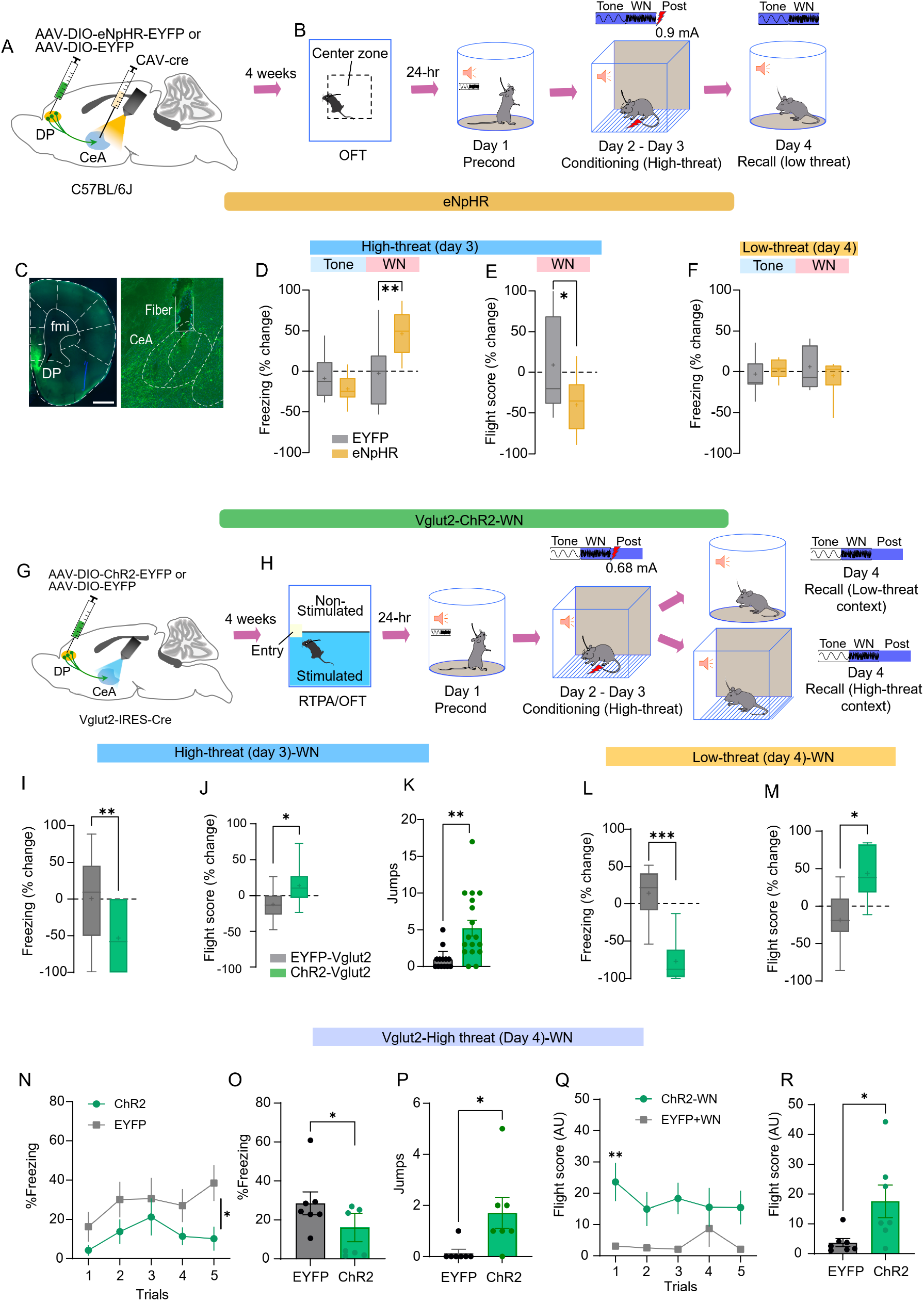
Optogenetic modulation of the DP-to-CeA pathway regulates flight. **A**, Schematic of intersectional targeting strategy for optogenetic inhibition of DP-to-CeA pathway. **B**, Experimental timeline for optogenetic inhibition experiments. **C**, Representative images of eNpHR-EYFP expression in DP and fibre stub placement in CeA (scalebar = 500 µm). **D**, Effect of optogenetic inhibition in EYFP (N = 9) and eNpHR (N = 9) groups on freezing (LED-on vs LED-off) in the high-threat context (EYFP vs eNpHR, Mann-Whitney, ** *p* < 0.01). **E**, Effect of optogenetic inhibition in EYFP (N = 9) and eNpHR (N = 9) groups on flight scores (LED-on vs LED-off) in the high-threat context (EYFP vs eNpHR, Mann-Whitney, * *p* < 0.05). **F**, Effect of optogenetic inhibition in EYFP (N = 9) and eNpHR (N = 9) groups on freezing (LED-on vs LED-off) in the low-threat context (EYFP vs eNpHR, Mann-Whitney, n.s.) **G**, Schematic of viral injection and strategy for optogenetic stimulation of DP-to-CeA pathway. **H**, Experimental timeline for optogenetic stimulation experiments. **I**, Effect of optogenetic stimulation in EYFP control (N = 13) or Vglut2-ChR2 (N = 17) groups on freezing (LED-on vs LED-off) in the high-threat context (Unpaired t-test, t=2.46, df=28, \**p* = 0.02). **J**, Effect of optogenetic stimulation in EYFP control (N = 13) or Vglut2-ChR2 (N = 17) groups on flight scores (LED-on vs LED-off) in the high-threat context (Unpaired t-test, t=2.262, df=28, \**p* = 0.0316). **K**, Effect of optogenetic stimulation in EYFP control (N = 13), and Vglut2-ChR2 (N = 17) groups on total escape jumps in the high-threat context (Unpaired t-test, t=3.383, df=28, \*\**p* = 0.0021). **L**, Effect of optogenetic stimulation in EYFP control (N = 6) and Vglut2-ChR2 (N = 10) groups on freezing (LED-on vs LED-off) in the low-threat context (Unpaired t-test, t=5.046, df=14, \*\*\**p* < 0.001). **M**, Effect of optogenetic stimulation in EYFP control (N = 6) and Vglut2-ChR2 (N = 10) groups on flight scores (LED-on vs LED-off) in the low-threat context (Unpaired t-test, t=2.845, df=14, *p < 0.05). **N**, Trial-wise freezing during optogenetic stimulation in EYFP control (N = 7) and Vglut2-ChR2 (N = 7) groups during recall in the high-threat context. (Two-way ANOVA, Group x trial, F_(4, 48)_ = 0.644, P=0.633, main effect of Group, F _(1,12)_ = 4.874, P=0.0475). **O**, Average freezing during optogenetic stimulation in EYFP control (N = 7) and Vglut2-ChR2 (N = 7) groups during recall in the high-threat context (Unpaired t-test, t=2.208, df=12, \**p* < 0.05). **P**, Number of escape jumps during optogenetic stimulation in EYFP control (N = 7) and Vglut2-ChR2 (N = 7) groups during recall in the high-threat context (Unpaired t-test, t=2.524, df=12, \**p* < 0.05) **Q**, Trial-wise flight score during optogenetic stimulation in Vglut2 EYFP control (N = 7) and Vglut2 ChR2 (N = 7) groups during recall in the high-threat context (Two-way ANOVA, Trial x Group, F_(4, 48)_ = 2.738, P=0.0393, main effect of Group, F _(1,12)_ = 6.06, P = 0.0299, followed by Bonferroni’s multiple comparisons test. \*\**p* < 0.01). **R**, Average flight score during optogenetic stimulation in Vglut2 EYFP control (N = 7) and Vglut2 ChR2 (N = 7) groups during recall in the high-threat context (Unpaired t-test, t=2.462, df=12, \**p* < 0.05). Box and whisker plots indicate median, interquartile range, and min. to max. of the distribution, crosses indicate means. Values in bar graphs and line graphs represented as mean ± s.e.m.

To study the sufficiency of the DP-to-CeA pathway in mediating flight, anxiety, and aversive behaviour, we performed optogenetic gain-of-function studies (**Fig. 4G–R; Figs. S5–S8**). Optogenetic activation of DP-to-CeA terminals using a non-cell type specific intersectional approach (**Fig. S5A**) did not elicit significant changes in negative valence behaviours (**Fig. S5B– J**). Stimulation of DP terminals in the CEA expressing ChR2 under the control of a CaMKIIa promoter (**Fig. S6A**) elicited a significant avoidance response (**Fig. S6B, C**); however, there was no significant effect on freezing and flight responses in either threat context (**Fig. S6D–G**).

The lack of effect of these manipulations on defensive behaviour could be due to functional heterogeneity in the DP-to-CEA pathway (**Fig. 1J**). Therefore, we separately targeted the Vglut1+ and Vglut2+ subpopulations and stimulated axon terminals in the CEA (**Fig. 4G; Figs. S7, S8**). Stimulation of Vglut1+ terminals induced real-time place preference (**Fig. S7B, C**), and a reduction in cue-induced freezing (**Fig. S7D, F**), but it had no effect on conditioned flight responses (**Fig. S7E, G**). Optogenetic activation of Vglut2+ terminals in the CeA (**Fig. S8A–B**), however, induced significant place avoidance (**Fig. S8C**), and a significant decrease in the number of centre entries in the OFT (**Fig. S8D, E**). Stimulation of Vglut2+ terminals also significantly reduced freezing (**Fig. 4I, L; Fig. S8F, I**), and significantly elevated flight responses in both contexts (**Fig. 4J, K, and M; Fig. S8G, H, and J**).

We next tested the effects of Vglut2+ terminal stimulation on flight responses in the high-threat context in absence of footshock (**Fig. 4N–R**). Optogenetic stimulation during WN significantly reduced freezing (**Fig. 4N, O**), and increased flight responses (**Fig. 4P–R**). Together, these data demonstrate that the DP-to-CeA pathway is necessary and sufficient for conditioned flight responses.

## DP neurons control CeM outputs

To investigate how DP neurons might impact flight behaviour via the CeM, we first asked whether DP-to-CeA projections differentially innervate defined cell types in the CeM that are known to mediate defensive behaviour, as in the CeL^2^ (**Fig. S9A–I**). There were many labelled terminals near both somatostatin (SOM)+ and corticotrophin releasing hormone (CRH)+ somata in the CeM (**Fig. S9B**), and DP fibre stimulation resulted in a strong induction of either a monophasic inward synaptic current or a biphasic inward/outward synaptic current in both SOM+ and CRH+ neurons in the CeM (**Fig. S9D, F, and I**), but not CeL (**Fig. S9E, H**). These optogenetically-evoked responses were blocked by bath application of the AMPA receptor antagonist DNQX (**Fig. S9G**). These findings confirm the presence of functional glutamatergic DP projections to the CeM but also suggest that this pathway controls flight behaviour via a mechanism distinct from that which was previously described in the CeL.

CeM output neurons can be classified by their action potential firing properties,^24–26^ with distinct subclasses influencing behavioural threat responding via their projections to known targets within the periaqueductal gray (PAG).^26–28^ In particular, burst firing and regular firing neurons innervate the dorsolateral and lateral columns of the PAG (dl/l PAG) involved in flight responses.^26,29,30^ We tested if DP innervation targeted physiologically defined CeM neuronal populations. We classified DP-excited CeM neurons as regular spiking, bursting, or late-firing (**Fig 5A–C; Fig. S9J**). Optogenetic stimulation of DP terminals in the CeA primarily evoked EPSCs in burst firing and regular firing neurons, while late-firing neurons were only rarely excited (**Fig. 5C**), suggesting that the DP excites CeM projections to PAG columns linked to flight behaviour.

**Figure 5.**
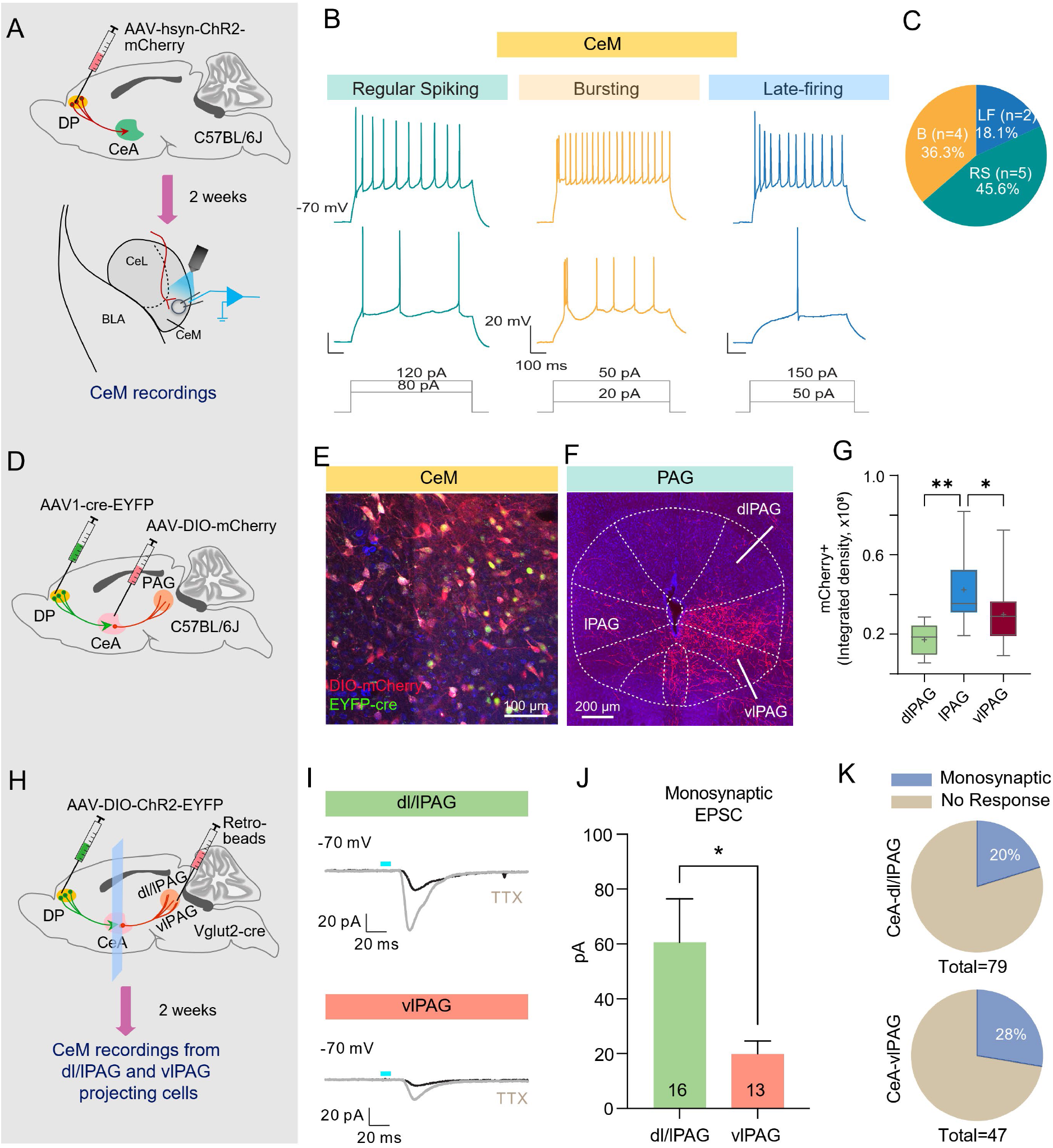
DP-to-CeA neurons exert excitatory control over CeM neurons projecting to PAG. **A**, Injection targeting and recording strategy. **B**, Representative evoked synaptic responses in CeM cells by photostimulation of DP axonal fibres and following DNQX application in voltage-clamp (-70 mV). **C**, Proportion of DP-excited CeM neurons classified by firing patterns (N = 11 cells from 4 mice). **D**, Schematic of viral targeting strategy to map the DP innervating CeM neurons projecting to PAG subregions. **E**, Representative image showing expression of EYFP-cre (green) and DIO-mcherry colocalized in the CeM. **F**, Representative image showing mcherry+ terminals in the PAG subregions. **G**, Morphometric analysis showing that lPAG receives significantly higher DP-CeM mCherry+ terminals (repeated measure One way-ANOVA, F_(2,36)_ = 12.50, P = 0.0006, followed by Bonferroni’s multiple comparison test, *p < 0.05, **p < 0.01). Box and whisker plots indicate median, interquartile range, and crosses indicate means. **H**, Schematic of targeting and recording strategy. **I**, Representative evoked synaptic responses in dl/lPAG (*top*) and vlPAG (*bottom*) projecting CeM cells by photostimulation of DP axonal fibres and TTX application. **J**, Average amplitude of evoked monosynaptic EPSCs was significantly higher in dl/lPAG (N = 16 cells from 10 mice) as compared to vlPAG (N = 13 cells from 7 mice) projecting CeM neurons (Mann Whitney test, *p = 0.0367). Values represented as mean ± s.e.m. **K**, Proportion CeM neurons with EPSCs by DP terminals stimulation, classified by their projections to dl/l PAG (20%, *top*) and vlPAG (28%, *bottom*).

To directly test this hypothesis, we first labelled the neuroanatomical pathway by injecting an anterograde transneuronal AAV1-Cre vector into the DP region and a Cre-dependent mCherry-expressing vector into the CeM (**Fig. 5D**). The projection topography of the labelled CeM neurons in the PAG revealed that the lateral PAG, known to regulate flight,^28^ had the highest density of mCherry+ fibres (**Fig. 5E–G**). We then labelled CeM projections to different columns of the PAG and performed whole cell patch clamp recordings while optogenetically stimulating Vglut2+ DP afferents in CeM (**Fig. 5H**). We observed that CeM projections to the dl/l PAG have larger monosynaptic excitatory currents than projections to the vlPAG in response to stimulation of DP terminals (**Fig. 5I–J**), despite there being no preferential innervation bias between the projection pathways (**Fig. 5K**). These data support the hypothesis that the DP promotes flight through excitation of CEA output neurons to columns of PAG known to be important for flight behaviour.

## Discussion

Here, we define the function of a novel top-down pathway from the DP region of the mPFC directly to the CeA in the control of defensive behaviour. This non-canonical corticolimbic pathway is distinct from the well-studied reciprocal pathway between the prelimbic and infralimbic cortices and the basolateral amygdala and is specifically activated in high-threat situations. Glutamatergic DP neurons exert their influence via projections to the CeM, the amygdalar output nucleus known for coordinating complex responses to threats. These findings have important implications for understanding the basic neurobiology of threat processing.

This adds to a growing body of literature defining the CeA as a vital limbic brain structure that integrates complex sensory information to generate survival behaviour.^23,31–33^ It is hypothesized that CeM output neurons control freezing through projections to the PAG,^28,34^ and our results demonstrate for the first time that the CeM is also important for flight. DP-to-CeA projections target specific classes of CeM neurons that project to flight centres in the PAG, with weaker inputs to neuronal classes linked to freezing.^26^ Our study supports the idea that top-down integration of information in the dl/l PAG contributes to defensive action selection to threat.^35,36^ It has been demonstrated that CeL circuit activity is regulated by local recurrent inhibitory interactions,^2,37,38^ and this mutual inhibition motif is likely important for rapid action selection. Future studies should investigate whether a similar “winner-take-all” motif is used in the CeM to select between freezing versus flight responses.

Learned fear is traditionally assayed using freezing, control of which involves the well-established mPFC-basolateral amygdala-CeA pathway.^39^ Previous research has shown that IL projections to the basolateral amygdala and intercalated cell masses effectively operate to inhibit fear by supporting fear extinction.^40,41^ In contrast, the DP-to-CeA pathway defined here operates under high-threat conditions and is necessary for generating flight. The context- and cue-specificity of DP-to-CeA activation and function is consistent with the known role of context and salience in defensive action selection.^42–44^ Our findings therefore implicate the importance of the mPFC in executive control over high-intensity fear responses and may lead to a better understanding of the cortical dysfunction observed in PTSD and panic disorder.^14^

## Methods

### Animals

Male and female C57BL/6J (Stock No: 000664, Jackson laboratories, USA) mice aged 8-12 weeks were used. CRH-IRES-Cre (Stock Number: 012704), SOM-IRES-Cre (Stock No: 013044), Vglut1-IRES-Cre (Slc17a7-IRES2-Cre-D, Stock No: 023527) and Vglut2-ires-cre (Stock Number: 028863) mice were used for *in-vitro* electrophysiology (aged 5-8 weeks), and to determine anatomical connectivity between DP and CeA, and CeA and PAG. For Vglut subpopulation tracing, Vglut1-IRES2-Cre and Vglut2-IRES-cre (Stock Number: 028863) mouse lines were used. All Cre-driver lines were fully backcrossed to C57BL/6J. All mice were individually housed on a 12-hr light/dark cycle with *ad libitum* food and water. Behavioural experiments were performed during the light phase. Animals were habituated by cupping the mouse in hand for 5 minutes each day for 2 days, before the start of the experiment. All animal procedures were performed in accordance with institutional guidelines and were approved by the Tulane University Institutional Animal Care & Use Committee.

### Viruses

For Cre-dependent expression of channelrhodopsin (ChR2), we used AAV5-EF1a-DIO-hChR2(H134R)-EYFP-WPRE-HGHpa (Addgene 20298, a gift from Karl Deisseroth). For Cre-dependent expression of halorhodopsin (eNpHR), we used AAV5-EF1a-DIO-eNpHR3.0-EYFP (Addgene 26966, a gift from Karl Deisseroth). Control mice for optogenetic experiments were injected with AAV5-EF1a-DIO-EYFP (Addgene 27056, a gift from Karl Deisseroth). For Cre-dependent expression of excitatory DREADDs, we used AAV5-hSyn-DIO-hM3D(Gq)-mCherry (Addgene 44361, a gift from Bryan Roth). For Cre-dependent expression of inhibitory DREADDs, we used AAV-hSyn-DIO-hM4D(Gi)-mCherry (Addgene 44362, a gift from Bryan Roth). For Cre-dependent GCaMP6f expression, we used AAV5-CAG-Flex-GcaMP6f-WPRE-SV40 (Addgene 100835, a gift from Douglas Kim & GENIE Project). To express Cre in C57BL/6J mice, we used retrograde transporting canine adenovirus type 2 carrying Cre-recombinase (CAV2-Cre, PVM, France).^45^ For anterograde neuronal tracing experiments in Vglut1- or Vglut2-cre mouse lines, AAV5-hSyn-DIO-mCherry (Addgene 50459, a gift from Bryan Roth) was injected in the DP.

To label CRH+ and SOM+ somata in the CeA for *in-vitro* electrophysiology experiments and anatomical tracing, CRH-IRES-Cre and SOM-IRES-Cre mouse lines were injected with AAV5-EF1a-DIO-EYFP (Addgene 27056, a gift from Karl Deisseroth). In the same mice, we used AAV5-hSyn-hChR2(H134R)-mCherry (Addgene 26976, a gift from Karl Deisseroth) to express ChR2 in the DP for *in-vitro* electrophysiology. To characterize firing patters of CeM neurons (unlabeled) innervated by DP projectors, we used AAV5-hSyn-hChR2(H134R)-mCherry (Addgene 26976, a gift from Karl Deisseroth) in DP. To study the direct effect of Vglut2+ DP projections of PAG projecting CeA neurons using *in-vitro* electrophysiology, we injected AAV5-EF1a-DIO-hChR2(H134R)-EYFP (Addgene 20298, a gift from Karl Deisseroth) in the DP of Vglut2-IRES-cre mice and retrograde beads in PAG subcolumns to label CeM projectors. For anterograde neuronal tracing experiments in Vglut1- or Vglut2-cre mouse lines, AAV5-hSyn-DIO-mCherry (Addgene 50459, a gift from Bryan Roth) was injected in the DP.

### Surgery

#### Tracers/Virus injections

Mice were deeply anaesthetized using 5% isoflurane (Fluriso, VetOne, Boise, ID) in oxygen-enriched air (OxyVet O2 Concentrator, Vetequip, Pleasanton, CA), followed by a subcutaneous injection of 2 mg/kg meloxicam (OstiLox, VetOne, Boise, ID), and then fixed into a stereotaxic frame (Model 1900, Kopf Instruments, Tujunga, CA) equipped with a robotic stereotaxic targeting system (Neurostar, Germany). Anesthetized mice were maintained on 2-2.5% isoflurane, and a core body temperature was maintained at 36°C using a feedback-controlled DC temperature controller (ATC2000, World Precision Instruments, Sarasota, FL). Eye ointment (GenTeal, Alcon, Switzerland) was applied to the mouse’s eyes to prevent dryness. The head was shaved, and the skin was sterilized using Betadine iodine solution (Purdue Products, Stamford, CT). 2% lidocaine (0.1 ml, Lidocaine 2%, VetOne, Boise, ID) was injected subcutaneously at the site of incision and a midline incision was made with a scalpel to expose the skull.

For the retrograde neuronal tracings, retrogradely transported beads (0.2 μl, 1:2 diluted with saline, Lumafluor Inc., Durham, NC) were stereotaxically injected into the respective brain regions. We injected CAV2-Cre into the CeA, and Cre-dependent AAV opsins or DREADD vectors into the DP. Approximately 0.4 μl per hemisphere of each vector was delivered bilaterally into the respective brain regions using pulled glass pipettes (tip diameter 10-20 μm, PC-100 puller, Narishige, Japan), connected to a pressure ejector (PDES-Pressure Application System, npi electronic equipment, Germany). AAV-DIO-mcherry was injected into the DP of Vglut1-/Vglut2-IRES-Cre mice for neuronal anterograde tracings. For *in-vitro* electrophysiology, AAV-syn-ChR2-mCherry was injected into the DP and Cre-dependent AAV-DIO-EYFP was injected into the CeA of CRH-IRES-Cre or SOM-IRES-Cre mice. AAV-DIO-ChR2-EYFP was injected into the DP of Vglut1-/Vglut2-IRES-Cre mice. For calcium imaging, C57BL/6J mice were injected unilaterally with CAV2-cre in the CeA and Cre-dependent GCaMP6f in the DP.

The CeA coordinates used were: 1.2-1.3 mm posterior to bregma, ±2.8-2.9 mm lateral to the midline and 4.1-4.3 mm below the dura. The DP coordinates used were 1.8-1.9 mm anterior to bregma, ±0.4-0.5 mm lateral to the midline and 3.1-3.2 mm below the dura. The DMH coordinates used were 1.7 mm posterior to bregma, ±0.5 mm lateral to the midline and 5 mm below the dura. The dl/l and vl PAG coordinates used were -4.16 mm (-4.72 mm for vlPAG) posterior to bregma, -0.4 mm lateral to the midline and 2.4 mm (2.7 mm for vlPAG) below the dura.

#### Fibre implantation for optogenetics

For optogenetic manipulations of behaviour, animals were bilaterally implanted with LC optic fibre stubs (fibre: 0.48 NA, 200-230 μm diameter, 6 mm length, Plexon) two-three weeks after viral vector injection. Optic fibre tips were lowered to 100–200 μm above the CeA. Implants were fixed to the skull with two miniature screws (00.90-100-M-SS-FH, US Micro Screw, Seattle, WA), cyanoacrylate glue gel (SuperGlue, The original super glue, Ontario, CA) and dental cement (Ortho-Jet powder and black liquid acrylate, Lang, Wheeling, IL). Experiments were conducted four weeks after viral infections, to ensure adequate opsin expression.

#### Lens implantation for Ca^2+^ Imaging

Gradient index (GRIN) lens implantation surgery was performed as described previously.^46,47^ Mice were head fixed on the stereotaxic apparatus and a craniotomy was drilled over the DP coordinates, using 1.2 mm diameter round drill burr (Harvard Apparatus, Holliston, MA). Three additional holes were drilled to implant stainless steel screws (US Micro Screw, 00.90-100-M-SS-FH). The skull surface was cleaned from bone fragments and wiped using sterilized cotton tips. We aspirated the overlying tissue up to 1-1.5 mm above the site of implantation with a bent 27G needle (NE-4527, Component supply, Sparta, TN) and performed intermittent sterile saline washes to reduce intracranial pressure and improve the quality of the imaging site. A GRIN lens (1 mm diameter, 9 mm length, 0.5 NA, 1 pitch, part # 1050-002177, Inscopix, Palo Alto, CA) was attached to the stereotaxic arm using a custom-built lens holder, and lowered for 3 mm over the course of 10 min. When the lens reached the required depth, Kwik-Sil (World Precision Instruments, Sarasota, FL) was used to fill in the gap between the lens and the skull and left to dry for 5 min. To hold the lens firmly in place, an adhesive cement (C&B S399 Metabond Quick Adhesive Cement System Parkell, Edgewood, NY) mixed with liquid polymer (Catalyst + Quick Base) was applied around the lens. After that, dental acrylic cement (Ortho-Jet powder and clear liquid acrylate) was applied over the skull and screws and allowed to dry completely. The lens holder was loosened and carefully detached from the lens. The layers of dental cement mixed with black acrylate (Ortho-Jet powder and black liquid acrylate) were applied around the lens until the height was ∼0.5 mm below the top of the lens, and a thick layer of Kwik-cast (World Precision Instruments, Sarasota, FL) was applied over the lens until it was embedded. This ridge around the lens prevented lens damage during homecage activity and was used later for baseplate implantation. Animals were then returned to the homecage, individually housed, and allowed to recover for 2-3 weeks.

#### Baseplating and verification of Ca^2+^ transients

Mice were checked weekly for GCaMP6f fluorescence and Ca^2+^ transient activity over several weeks after the lens implantation, as described before.^47^ For baseplating, mice were anesthetized with 5% isoflurane, fitted into the stereotaxic frame, and then maintained at 1%. A head-mounted miniscope (V4, UCLA open-source, http://miniscope.org/) was fixed on a custom-built holder that was attached to the stereotaxic arm.^48^ The focusing mechanism of the miniscope was set at mid-range to leave enough distance for focus adjustments in either direction. The coaxial cable from the miniscope was attached to the data acquisition box (DAQ-V3.2, UCLA), which was connected to the computer through USB 3.0. The baseplate was attached to the bottom of the miniscope. We aligned the miniscope objective lens over the relay lens implanted in the mouse brain, and the miniscope was moved to approach the relay lens. The recording software (Miniscope controller, UCLA) was turned on and LED light intensity (25%), gain, and frame rate (30 fps) were set. To ensure maximum field of view and focus on the centre of the lens, the miniscope tilt was adjusted to view all edges of the relay lens. Then, the miniscope was slowly moved in X, Y, Z planes to locate active cells/transients. If there were no transients, the mouse was returned to the homecage and checked again the following week.

If Ca^2+^ transients were observed, they were further confirmed using tail pinch. If multiple cells were observed, these mice were then considered for baseplating. After finding the optimal focal plane and clear field of view, the baseplate was fixed on the mouse skull. Dental cement mixed with black acrylate (Ortho-Jet powder and black liquid acrylate) was applied layer-by-layer between the baseplate and the circular ridge built during lens implantation. The dental cement was applied sufficiently so that no light could pass through from the outside. After the cement was dry, the miniscope was detached and a plastic cover was installed to protect the relay lens.

### Miniscope recordings

Animals with Ca^2+^ transients were used for behavioural experiments (N = 6). First, the mice were subjected to preconditioning on day 1, and their behaviour as well as Ca^2+^ responses were recorded. Animals were conditioned in the high-threat context for 2 days (5 trials/day, 0.9 mA shock intensity), followed by recall in the low-threat context on day 4 (4 trials/day). The Ca^2+^ response data was at 30 frames/sec rate. Behaviour was simultaneously recorded to video (“Pike” camera, Allied vision, Germany). Neuronal recordings were started 10 s before the start of each trial (at the onset of preSCS) and turned off 10 sec after the end of shock/ cue (post-shock/cue), while behaviour was continuously recorded. Mice with flight responses only included in the for the ca^2+^ data analysis.

### Ca^2+^ imaging processing

Ca^2+^ imaging videos were concatenated using ImageJ software (https://imagej.nih.gov/ij/download.html). The concatenated videos were then processed using MATLAB and an open source analysis package (https://bahanonu.github.io/ciatah/) as described previously.^47,49^ The videos were pre-processed by correcting for motion and temporally downsampled by a factor of six. After pre-processing the videos, we extracted individual neurons and their activity traces by using the open-source package Constrained Nonnegative Matrix Factorization for microEndoscopic data (CNMF-E; https://github.com/zhoupc/CNMF-E). The extracted cells were then manually identified based on the spatial filter and activity trace of each candidate cell along with the candidate cells’ average Ca^2+^ transient waveform. Data were averaged over a 200 ms sliding window. The data was normalized by calculating Z-scores (subtracting mean activity scores during all preSCS trials from each activity trace, divided by the standard deviation of each cell during all preSCS trials) and the neurons with peaks that were ≥3 s.d. above baseline were considered as active neurons. To determine statistically significant changes in responses of the DP-to-CeA projecting cells to the tone and WN, the Z-scores from the corresponding periods were averaged and compared.

After finishing the experiments animals were perfused and their brains were isolated to confirm the lens placements. The data from animals with correct lens placement was used for analysis.

### Behavioural paradigms

#### Conditioned flight paradigm

Two different contexts were used. Context A (low-threat context) consisted of a clear cylindrical chamber with a smooth floor, while Context B (high-threat context) consisted of a square enclosure with an electrical grid floor used to deliver alternating current footshocks (ENV-414S, Med Associates Inc., Fairfax, VT). These two chambers were cleaned with 1 % acetic acid and 70 % ethanol, respectively. A speaker (ENV-224AM, Med Associates Inc.) was mounted above the chambers to deliver auditory stimuli at 75 dB. A programmable audio generator (ANL-926, Med Associates, Inc.) generated auditory stimuli. Behavioural protocols were generated using MedPC software (Med Associates, Inc.) to control auditory stimuli and footshock via TTL pulses.

We used a four-day paradigm as described previously.^2,21^ On Day 1, animals were subjected to pre-exposure in context A. After a 3 min baseline period, there were 4 presentations of a serial compound stimulus (SCS, 20 s long) with a 90 s average pseudorandom intertrial interval (ITI) (range 80−100 s). The SCS consisted of a 10 s tone (7.5 kHz, 500 ms pips at 1 Hz) and a 10 s white noise (WN, random distribution of 1 to 20,000 Hz, 500 ms pips at 1 Hz). The total duration of the pre-exposure session was 590 s. On Day 2 and Day 3 (conditioning days), mice were placed in Context B and, after a 3 min baseline period, presented with five pairings of the SCS with a 120 s average pseudorandom ITI (range 90−150 s) co-terminating with a 1 s, 0.9 mA footshock (0.68 mA shock for ChR2 mediated stimulation groups). Each conditioning session lasted for 820 s. Recall session on day 4 was conducted in Context A (or Context B for recall in high-threat context group). Following a 3 min baseline period, mice were presented with 4 trials of the SCS without footshock, with a 90 s average pseudorandom ITI (range 80−100 s) over a total period of 590 s.

For optogenetic inhibition/control experiments the mice were presented with 8 trials on Day 3 with 4 pairs of alternating light ON-OFF trials (counter-balanced). During recall mice were presented with 3 pairs of alternating light ON-OFF trials.

For rest of the optogenetic excitation/control experiments the mice were presented with 5 trials on Day 3 with 2 pairs of alternating light ON-OFF trials (counter-balanced). During recall mice were presented with 2 pairs of alternating light ON-OFF trials (counter-balanced). For recall experiment in the high-threat context, we presented mice with 5 trials, all ON, for ChR2 and EYFP controls.

#### Quantification of defensive behaviour

All sessions were recorded to video, and behaviour was analysed using contour tracking (Cineplex software, Plexon, Dallas, TX). Freezing was defined as a complete cessation of movement for at least 1 s and was automatically scored using a frame-by-frame analysis of pixel changes (Cineplex Editor, Plexon). Results were confirmed by a trained observer blinded to condition. By determining a calibration coefficient using the chambers’ known size and the camera’s pixel dimensions, speed (cm/s) was extracted using the animal’s centre of gravity.^21^

Escape jumping was scored manually from video files by a blinded observer. Flight scores were calculated by dividing the average speed during each CS by the average speed during the 10 s pre-CS from all the trials (baseline, BL) and then adding 1 point for each escape jump (speedCS/ speedBL + # of jumps). A flight score of 1 therefore indicated no change in flight behaviour from the preSCS period.

#### Open field test

A 45 × 45 × 45 cm arena made of white opaque Plexiglas was used. The arena was cleaned using 70% ethanol after each mouse. Mouse behaviour was tracked under 100 lux light conditions using a top-mounted camera (“Pike” camera, Allied vision, Germany).

*For chemogenetic* experiments, the OFT duration was 10 min.

*For optogenetic experiments*, the OFT duration was 6 min. After a 120 s baseline period, 4, 30 sec ON-OFF trials of light stimulation were used. For eNpHR experiments, a continuous orange light was used at 10 mW power at the fibre tip, and for ChR2 experiments, 10 ms pulses of blue light were used at 10 Hz, and 10 mW power at the fibre tip.

The time spent in the inner zone, the number of entries into it, and total distance traveled were measured using tracking software (Cineplex Studio, Plexon).

#### Elevated plus maze

Behaviour was tracked using a top-mounted camera (“Pike” camera, Allied vision, Germany) for 10 min. The maze was made from white opaque Plexiglas material and consisted of four 30 cm long and 7 cm wide arms. Two open arms had no walls, and two enclosed arms had 15 cm high walls. The maze was elevated 40 cm above the ground. The arena was cleaned using 70% ethanol after every test. The total time spent in the open arms was calculated using Cineplex Editor software (Plexon).

### Chemogenetic manipulations of behaviour

Behavioural testing was performed 3-4 weeks after viral injections. First, mice were subjected to the OFT. On the next day, animals were subjected to the EPM. Thirty min before the start of behavioural testing animals were injected with clozapine N-oxide [CNO; 0.5 mg/ml in vehicle (0.9% saline), given as 10 ml/kg for final dose of 5 mg/kg, intraperitoneally (ip); Enzo Life Sciences, Farmingdale, NY] or vehicle (10 ml/kg volume, ip).

### Optogenetic manipulations of behaviour

Two-three weeks after vector injection, animals were implanted with fibre stubs (Plexon) targeting CeA, as described in Surgery. For optogenetic modulation, we used a Plex controller system (Plexon) operated through Radiant software (Plexon). Two PlexBright LED modules (Plexon) were connected to the controller, and an LC patch cable (200-230 µm fibre, 1 meter long, 0.66 NA, Plexon) was connected. Laser power at the fibre tip was measured before every test with an optical power and energy meter (PM100D, ThorLabs, Newton, NJ). The patch cable was connected to the head-mounted fibre stubs using ceramic sleeves. Connectors were tested for coupling efficiency before implantations, and laser power at the fibre tip for behavioural manipulations was adjusted to reach a value of 10 mW.

Experiments were performed with a within-subjects design in which mice underwent eight trials (5 trials for ChR2 groups) during FC2 with alternating 2-2 trials of LED ON-OFF. During recall session, mice underwent six trials (4 trials for ChR2 groups) with alternating light on off conditions. To inhibit (eNpHR) DP-CeA neuronal populations during threat cues, a 620 nm light was switched on 500 ms before the onset of the SCS (first tone pip) and remained on until the end of the last white noise pip or (20.5 s in total).

For ChR2-induced excitation (0.9 mA footshock intensity group) of DP-to-CeA neuronal populations, we used 465 nm light pulses (10 Hz, 10 ms width) which was switched on 500 ms before the onset of the SCS (first tone pip) and remained on until the end of the last white noise pip (20.5 s in total).

In addition, for ChR2 manipulation during low footshock intensity (0.68 mA) groups, the 465 nm light pulses (10 Hz, 10 ms width) were delivered when first WN pip was on and remained on until the end of 10 sec post cue periods (20.5 s in total).

### Real-time place aversion test

To test whether optogenetic stimulation of the DP-to-CeA pathway induced aversive responses, we subjected the AAV-DIO-ChR2-EYFP or AAV-DIO-EYFP-injected mice to real-time place aversion test for 10 min. We used a custom-made 50 × 50 × 50 cm arena made from white opaque Plexiglas and divided into two equal compartments. One side of the chamber was allotted as the stimulation side and other as a neutral side (non-stimulated, with striped walls and metallic plate floor). Mice were placed in the non-stimulated side, and 10 Hz blue LED light stimulation was delivered each time the mouse moved to the stimulation side of the chamber. The light stimulation was continuously on until the mouse moved back into the non-stimulation compartment. Behaviour was recorded via a top mounted recording camera attached to recording system (Plexon) and scored using Cineplex editor (Plexon) by a blinded experimenter. The % time spent in the stimulation chamber was used as a measure of aversion.

### Immunohistochemistry (IHC)

Following the completion of experiments, mice were anesthetized with tribromoethanol (240 mg/kg) and perfused transcardially with phosphate-buffered saline (PBS) followed by 4% paraformaldehyde in PBS. Brains were isolated and stored in paraformaldehyde overnight. On the next day, fixed brains were sectioned using a vibrating microtome (Precisionary, Greenville, NC) in 60-(for cFos IHC) or 80-μm thick coronal slices.

Antibody staining was performed on floating tissue sections. Briefly, sections were washed in PBS-TritonX100 (PBST, 0.3%) and blocked using 5% goat serum in PBST for 1 hr followed by an overnight (for RFP or GFP staining), or 48-h (for cFos staining), incubation in primary antibodies at 4°C. Primary antibodies used in this study were rabbit anti-RFP (1:1500, 600-401-379, Rockland Immunochemicals, Pottstown, PA), chicken anti-GFP (1:2000, NB100-1614, Novus Biologicals, USA), rabbit anti-cFos (1:1000, 226003, Synaptic Systems, Germany). After primary antibody incubation, sections were washed in PBST and incubated in secondary antibodies in PBST (1:500 goat anti-rabbit AlexaFluor 555/488 or goat anti-chicken AlexaFluor 488, Thermo Fisher Scientific, USA). Following final rinses with PBS, sections were mounted and scanned from the DP and CeA.

Images were obtained using an Axio Scan.Z1 slide-scanning microscope (Zeiss, Germany) and a confocal microscope (FV3000, Olympus, Japan). Mice were included in subsequent data analyses only if bilateral expression specific to the target region was observed.

### cFos quantification

Red fluorescent retro-beads were injected into the CeA of C57BL/6J mice. 7 days later, animals were divided into 4 groups: homecage control, shock-only, SCS-FC and unpaired. Shock-only, unpaired and SCS-FC mice were subjected to two days of conditioning and were sacrificed 90 min after the second session by transcardial perfusion. Following brain sectioning and IHC staining for cFos, confocal images were taken. The quantification of cFos+, bead+, and cells with both markers were performed by a blinded observer using ImageJ.

### Brain slice electrophysiology

#### Slice preparation

Coronal brain slices containing the CeA were collected from mice at least two weeks after viral injections for *ex vivo* electrophysiological recordings, as described earlier.^38,50^ Mice (9+ weeks) were decapitated in a restraining plastic cone (DecapiCone, Braintree Scientific, Braintree, MA) and the brains were dissected and immersed in ice-cold, oxygenated cutting solution containing the following (in mM): 252 sucrose, 2.5 KCl, 26 NaHCO_3_, 1 CaCl_2_, 5 MgCl_2_, 1.25 NaH_2_PO_4_, 10 glucose. The brains were trimmed and glued to the chuck of a Leica VT-1200 vibratome (Leica Microsystems, Germany) and 300 µm-thick coronal slices were sectioned. Slices were transferred to a holding chamber containing oxygenated recording artificial cerebrospinal fluid (aCSF) containing (in mM): 126 NaCl, 2.5 KCl, 1.25 NaH_2_PO_4_, 1.3 MgCl_2_, 2.5 CaCl_2_, 26 NaHCO_3_, and 10 glucose. They were maintained in the holding chamber at 34°C for 30 min before decreasing the chamber temperature to ∼20°C.

#### Patch clamp recording

Slices were bisected down the midline and hemi-slices were transferred from the holding chamber to a submerged recording chamber mounted on the fixed stage of an Olympus BX51WI fluorescence microscope equipped with differential interference contrast (DIC) illumination. The slices in the recording chamber were continuously perfused at a rate of 2 ml/min with recording aCSF maintained at 32-34°C and continuously aerated with 95% O_2_/5% CO_2_. Whole-cell patch clamp recordings were performed in eYFP-labeled SOM+ or CRH+ neurons, retrolabeled neurons (beads from dl/l or vl PAG), or unlabeled neurons in the CeM or CeL. Glass pipettes with a resistance of 3-5 MΩ were pulled from borosilicate glass (ID 1.2mm, OD 1.65mm) on a horizontal puller (Sutter P-97) and filled with an intracellular patch solution containing (in mM): 130 potassium gluconate, 10 HEPES, 10 phosphocreatine Na_2_, 4 Mg-ATP, 0.4 Na-GTP, 5 KCl, 0.6 EGTA; pH was adjusted to 7.25 with KOH and the solution had a final osmolarity of ∼ 290 mOsm. DNQX (20 uM) was delivered via the perfusion bath. Series resistance was normally below 15 MΩ immediately after break-in and was continuously monitored. Cells were discarded when the series resistance exceeded 30 MΩ. An optical fibre was placed approximately 2 mm above the slice and ChR2-expressing DP fibres were stimulated by 465 nm LED illumination (single 10-ms pulses,15 mW power, delivered at 1 Hz).

To assess firing properties in CeA neurons that displayed fast synaptic responses, 500-750 ms depolarizing current injections were applied in current clamp mode. To compare the properties of monosynaptic EPSCs evoked by optogenetic stimulation of DP fibers, responses were recorded in the presence of 1 mM 4-aminopyridine and 1 uM tetrodotoxin. Data were acquired using a Multiclamp 700B amplifier, a Digidata 1440A analog/digital interface, and pClamp 10 software (Molecular Devices, San Jose, CA). Recordings were filtered at 4 KHz and sampled at 50 KHZ. Data were analyzed with Clampfit 10 (Molecular Devices, San Jose, CA). Statistical comparisons were conducted with a paired or unpaired Students’s *t* test (p < 0.05 with a two-tailed analysis was considered significant).

### Statistical analysis

Data were analysed for statistical significance using Prism 10 (GraphPad Software, San Diego, CA). Statistical significance was set at *p* < 0.05. Data were tested for normal distribution using the Shapiro-Wilk normality test (α=0.05). For pairwise comparisons, the appropriate parametric (unpaired Student’s t-test) or nonparametric (Mann-Whitney test) test was performed. Data with more than two study groups were analysed using one-way ANOVA followed by Bonferroni’s post-hoc tests. For calcium imaging data analysis, we used custom code written in MATLAB. Correlations analysis and plotting regression lines were performed using the GraphPad Prism 10. The sample size was determined based on previously published research.

## Acknowledgements

We thank Dr. Biafra Ahanonu (University of California, San Francisco) for providing MATLAB codes and assistance with calcium imaging data analysis. We thank Dr. Ricardo Mostany (Tulane University, New Orleans) and Dr. Seongsik Yun (Northwestern University, Chicago) for help in standardization of calcium imaging and data analysis. This work was supported by the Louisiana Board of Regents through the Board of Regents support fund (**LEQSF(2018-21)-RD-A-17**) and *the National Institute of Mental Health* of the National Institutes of Health under award number **R01MH122561**. The content is solely the responsibility of the authors and does not necessarily represent the official views of the National Institutes of Health.

## Author contributions

Conceptualization— CDB, JPF; Formal analysis— CDB, CS, XF, MD, QEL, RV, CV, AW, SB, AD, EB, AR, JP, JPF; Funding acquisition— JPF; Investigation— CDB, CS, XF, QEL; Methodology— CDB, MD, CS, JT, JPF; Project Administration / Supervision— CDB, JPF, JT Resources— JPF, JT; Visualization— CDB, CS, MD, XF, JPF; Writing, original draft— CDB, JPF; Writing, review & editing— CDB, CS, XF, JT, JP, JPF

## Competing Interest Statement

The authors declare no competing interests.

## Additional Information

Correspondence and requests for materials should be addressed to JPF.

## Data availability

Data sets included in this study are available from the corresponding author upon reasonable request.

## Extended Data

**Figure S1.**
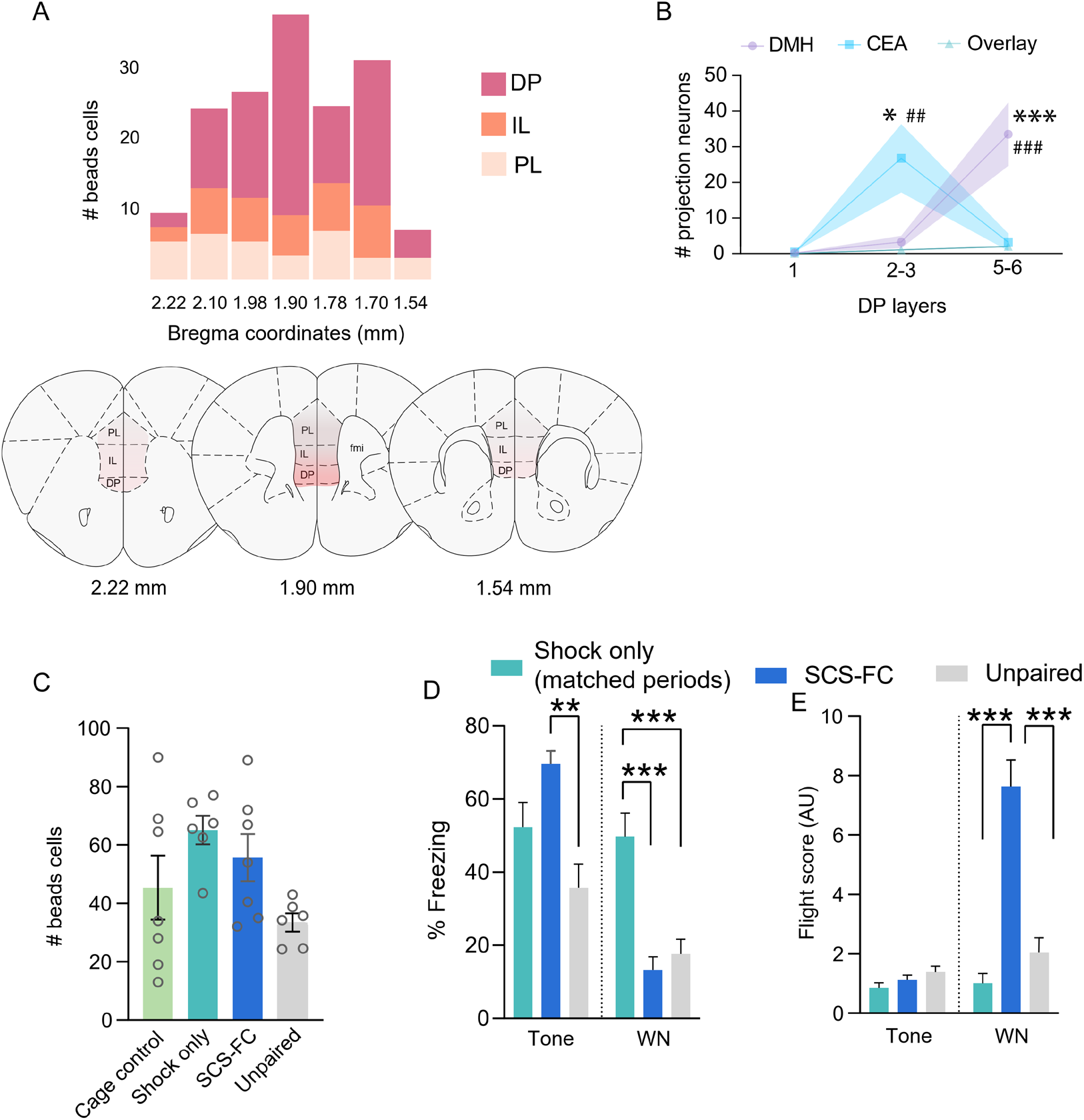
(Data related to Figure 1): Neuroanatomy of the DP-CeA pathway. **A**, *Top*, Number of CeA-projecting mPFC cells across the antero-posterior axis (2.22 mm to 1.54 mm). *Bottom*, Schematic of coronal sections showing the density of beads in DP on anterio-posterior scale. **B**, The layer-wise distribution of bead+ cells in the DP that project to CeA and/or DMH (N = 6; Two-way ANOVA, layer x group, F_(4, 45)_ = 10.15, *p* < 0.0001, followed by Bonferroni’s post-hoc test). \**p*<0.05, *** *p* <0.001 (DMH vs CeA), ^##^*p* <0.01, ^###^*p* <0.001 (vs overlay). **C**, Total number of bead+ cells across groups (N = 6; 3-4 slices per group). One-way ANOVA, F _(3, 22)_ = 2.819, *p* = 0.0626. **D**, Freezing of cFos groups on FC2. One-way ANOVA for tone F_(2, 15)_ = 8.502, *p* = 0.0034 and white noise (WN; F_(2, 15)_ = 18.43, *p* < 0.0001), followed by Bonferroni’s post-hoc test. \*\**p* < 0.01, *** *p* <0.001. **E**, Flight scores of cFos groups on FC2. One-way ANOVA for tone (F_(2, 15)_ = 1.87, *p* = 0.18) and white noise (WN) F_(2, 15)_ = 33.11, *p* < 0.0001, followed by Bonferroni’s post-hoc test. *** *p* <0.001.

**Figure S2.**
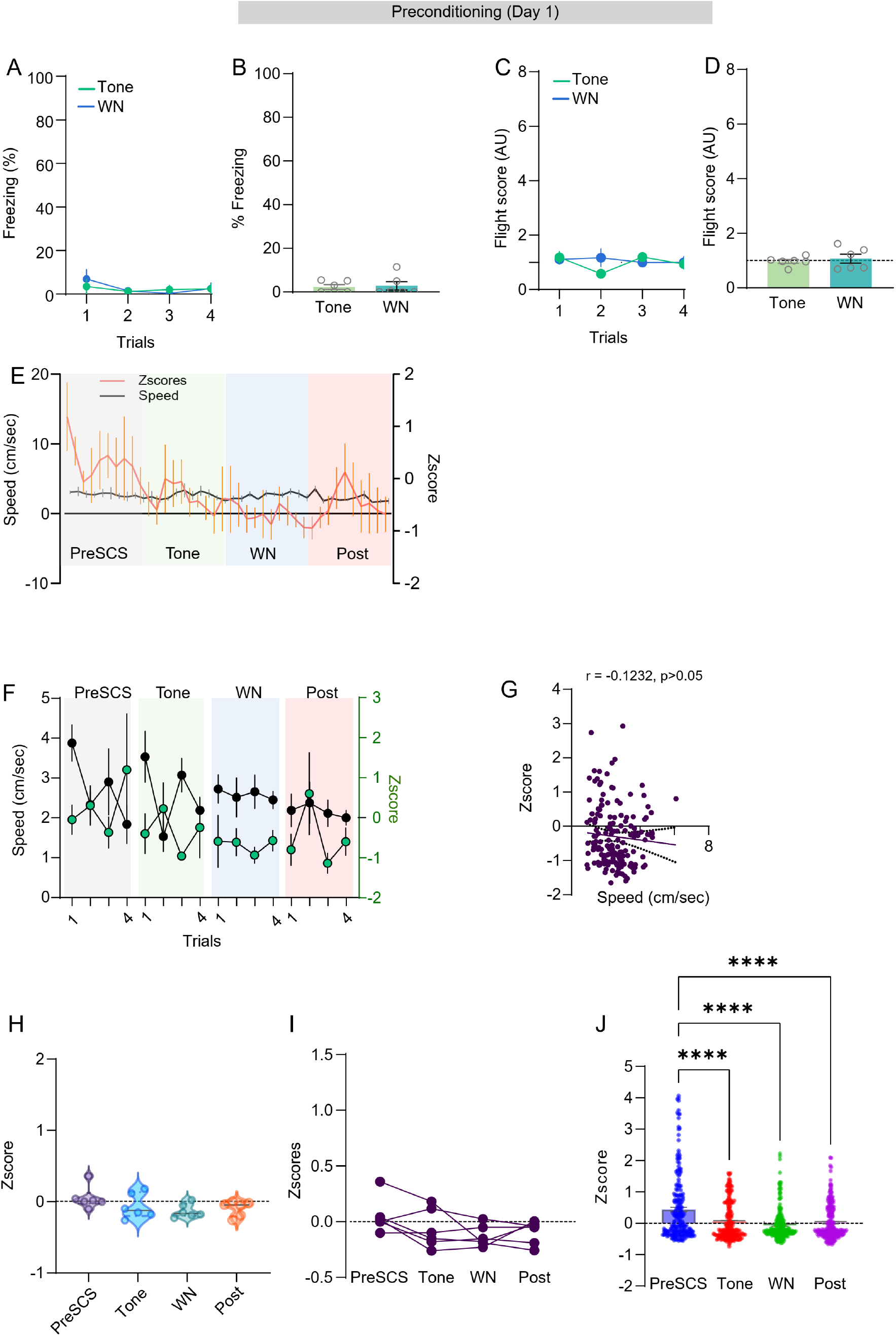
(Data related to Figure 2): Calcium imaging during pre-conditioning. **A-B**, Trial-wise and cumulative freezing of mice from calcium imaging experiments during preconditioning session (N = 6, Paired t-test, t=0.3051, df=5, p = 0.77). **C-D**, Trial-wise and cumulative flight score of mice from calcium imaging experiments during preconditioning session (N = 6, Paired t-test, t=0.6565, df=5, p = 0.54). **E**, Speed and neuronal activity during the last trial of preconditioning session (Day 1, n = 221)). **F**, Average speed and neuronal activity during each trial of preSCS, tone, WN and post-cue periods. **G**, Correlation of speed and neuronal activity from all trials (10 sec each epoch of preSCS, tone, WN and post cue, each point represents one sec). Spearman correlation, r = -0.1232, 95% CI: - 0.2774 to 0.03724, p >0.05. **H**, Average Z-score of the DP-to-CeA population during the preSCS, tone, WN and post-cue periods (Ordinary one-way ANOVA, F_(3, 20)_ = 1.965, p = 0.15). **I**, Average Z-scores of individual mice during preSCS, tone, WN and post-cue periods (N = 6). J, Z-scores of individual neurons during the last trial of preconditioning (N = 6, total neurons = 221, One-way ANOVA, F_(3, 880)_ = 21.43, P<0.0001, followed by Bonferroni’s multiple comparisons test, ****p<0.0001)

**Figure S3.**
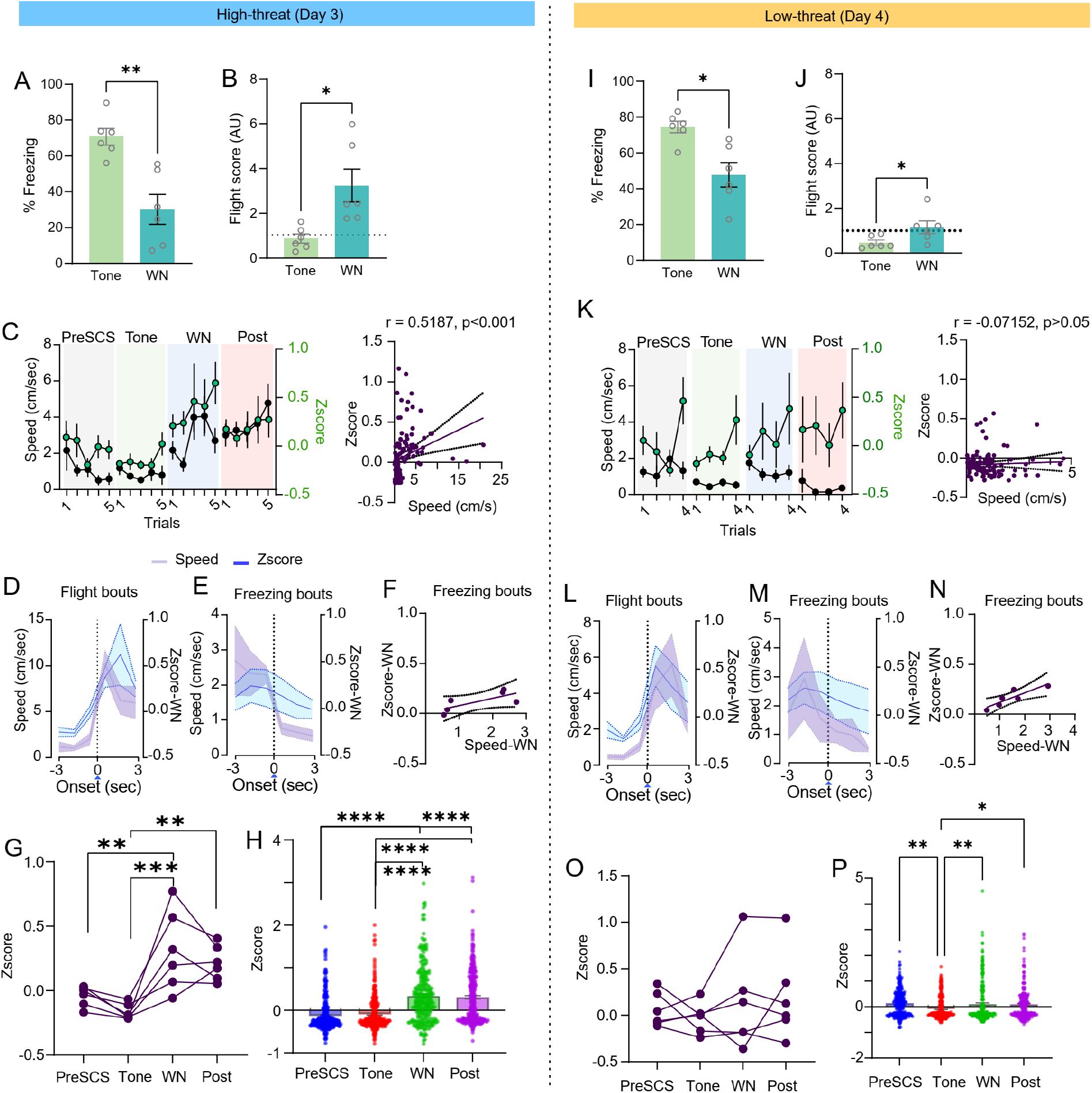
(Data related to Figure 2): Calcium imaging in the high-threat and low-threat contexts. **A**, Freezing behaviour in the high-threat context (N = 6). Paired t-test, t=4.744, df=5, \*\**p* < 0.01. **B**, Flight scores in the high-threat context (N = 6). Paired t-test, t=3.650, df=5, \**p* < 0.05. **C**, *left*, Average speed and neuronal activity during each trial of the preSCS, tone, WN and post-cue periods in the high-threat context (n = 273). Data shown as means ± s.e.m. *right*, Correlation of speed and neuronal activity from the last 3 trials (preSCS, tone, WN and post-cue epochs, each point represents data from 1 sec). Spearman correlation, r = 0.5187, 95% CI: 0.3696 to 0.6417, p<0.001. **D**, Speed and neuronal activity aligned to the onset of flight bouts during WN in the high-threat context. Data shown as means ± s.e.m. **E**, Speed and neuronal activity aligned to the onset of freezing bouts during WN in the high-threat context. Data shown as means ± s.e.m. **F**, Spearman correlation plot for speed and Z-score from the identified freezing bouts (each dot represents values at each sec of the bouts, r = 0.657, 95% CI = -0.02019 to 0.1662, p = 0.175). **G**, The linegraph showing the Z-scores of individual mice during preSCS, tone, WN and post-cue periods, across all trials (N = 6, One-way ANOVA, F_(3, 20)_ = 9.331, P=0.0005, followed by Bonferroni’s multiple comparisons test. \**p* < 0.05, \*\**p* < 0.01, ***p < 0.001). **H**, The Z-scores of individual neurons during preSCS, tone, WN and post-cue periods, from the last trial in the high-threat context (N = 6, total neurons = 273, One-way ANOVA, F_(3, 1112)_ = 59.01, P<0.0001, followed by Bonferroni’s multiple comparisons test, ****p < 0.0001). **I**, Freezing in the low-threat context (N = 6). Paired t-test, t=3.424, df=5, *p < 0.05. **J**, Flight scores in the low-threat context (N = 6). Paired t-test, t=2.889, df=5, *p < 0.05. **K**, *left*, Change in average speed and neuronal activity during preSCS, tone, WN and post-cue periods in the low-threat context (Day 4) over 4 trials (n = 253). Data shown as means ± s.e.m. *right*, Correlation of speed and neuronal activity from all recall trials in the low-threat context (preSCS, tone, WN and post cue epochs, each point represents 1 sec of data). Spearman correlation, r = -0.07152, 95% CI: -0.2526 to 0.1144, p>0.05. **L**, Speed and neuronal activity aligned to the onset of flight bouts during WN in the low-threat context. Data shown as means ± s.e.m. **M**, Speed and neuronal activity aligned to the onset of freezing bouts during WN in the low-threat context. Data shown as means ± s.e.m. **N**, Spearman correlation of speed and neuronal activity from freezing bouts (each point represents one sec of data, r = 0.82, 95% CI = 0.02337 to 0.1669, P = 0.058). **O**, Population activity of individual mice during preSCS, tone, WN and post-cue periods, across all trials (N = 6, One-way ANOVA, F_(3,20)_ = 0.3923, *P* = 0.75, followed by Bonferroni’s multiple comparisons test). **P**, Neuronal activity of individual neurons during preSCS, tone, WN and post-cue periods, from the last trial in the low-threat context (N = 6, total neurons = 253, One-way ANOVA, F_(3,1008)_ = 5.566, *P* = 0.0009, followed by Bonferroni’s multiple comparisons test, \**p* < 0.05, \*\**p* < 0.01).

**Figure S4.**
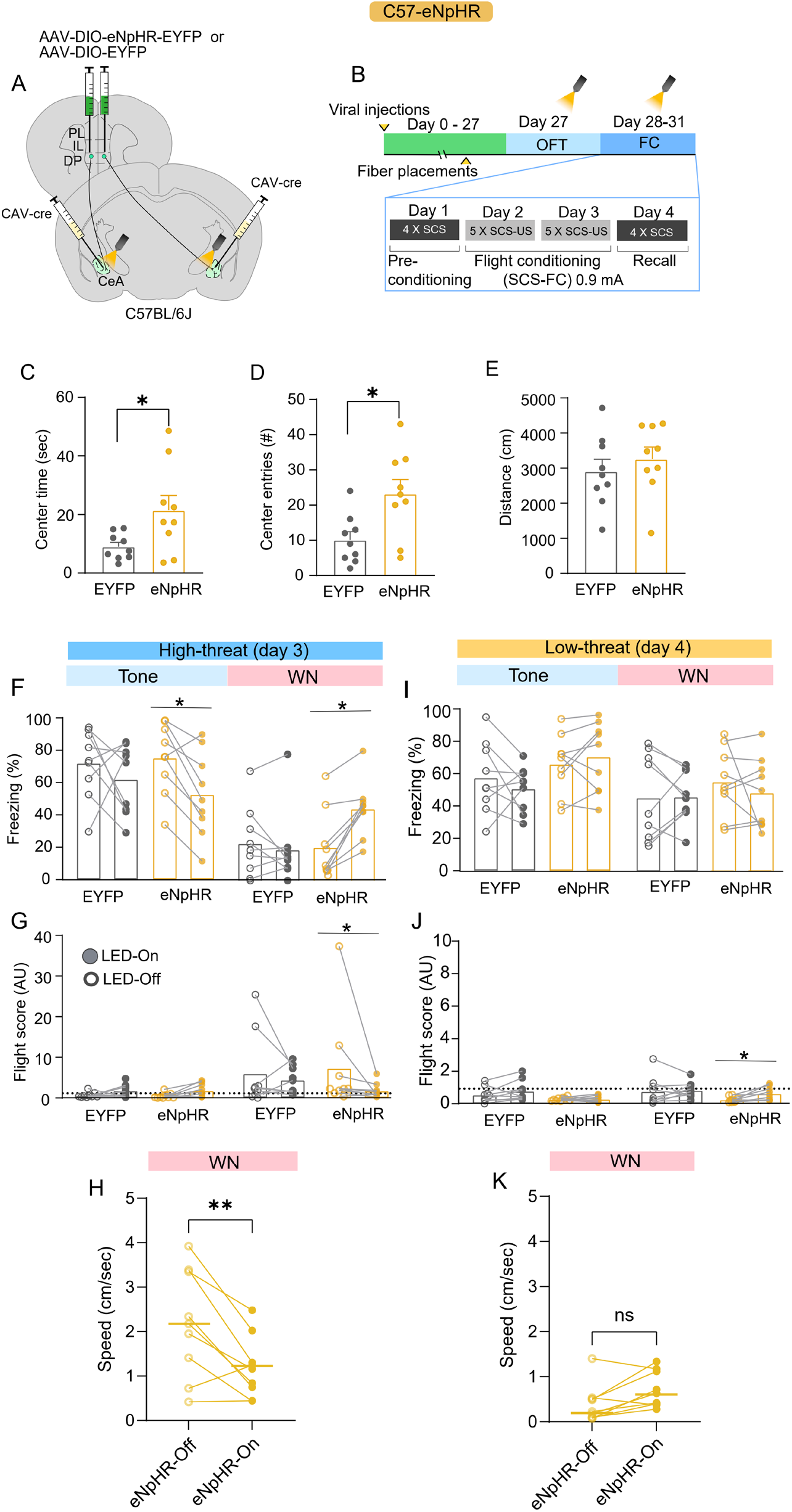
(Data related figure 4): Optogenetic inhibition of the DP-CEA pathway. **A**, Intersectional approach used for optogenetic terminal inhibition of the DP-to-CeA neuronal projections. **B**, Experimental timeline. **C-E**, Effect of optogenetic inhibition on centre time (**C**), centre entries (**D**), and distance travelled (**E**) in OFT in EYFP (N = 9) and eNpHR (N = 9) groups. Unpaired t-test, \**p* < 0.05. **F**, Effect of optogenetic inhibition in EYFP (N = 9) and eNpHR (N = 9) groups on freezing in the high-threat context (LED-on vs LED-off, Mann-Whitney, *P<0.05). **G**, Effect of optogenetic inhibition in EYFP (N = 9) and eNpHR (N = 9) groups on flight in the high-threat context (LED-on vs LED-off, Mann-Whitney, *P<0.05). **H**, Effects of optogenetic inhibition on speed in the high-threat context during WN in the eNpHR group (N = 9; Paired t-test t=3.497, df=8, p = 0.0081). **I**, Effect of optogenetic inhibition in EYFP (N = 9) and eNpHR (N = 9) groups on freezing in the low-threat context (LED-on vs LED-off, Mann-Whitney, n.s.). **J**, Effect of optogenetic inhibition in EYFP (N = 9) and eNpHR (N = 9) groups on flight in the low-threat context (LED-on vs LED-off, Mann-Whitney, *P<0.05). **K**, Effect of optogenetic inhibition on speed in the low-threat context during WN in the eNpHR group (N = 9; paired t-test,t=2.619, df=8, p = 0.307).

**Figure S5.**
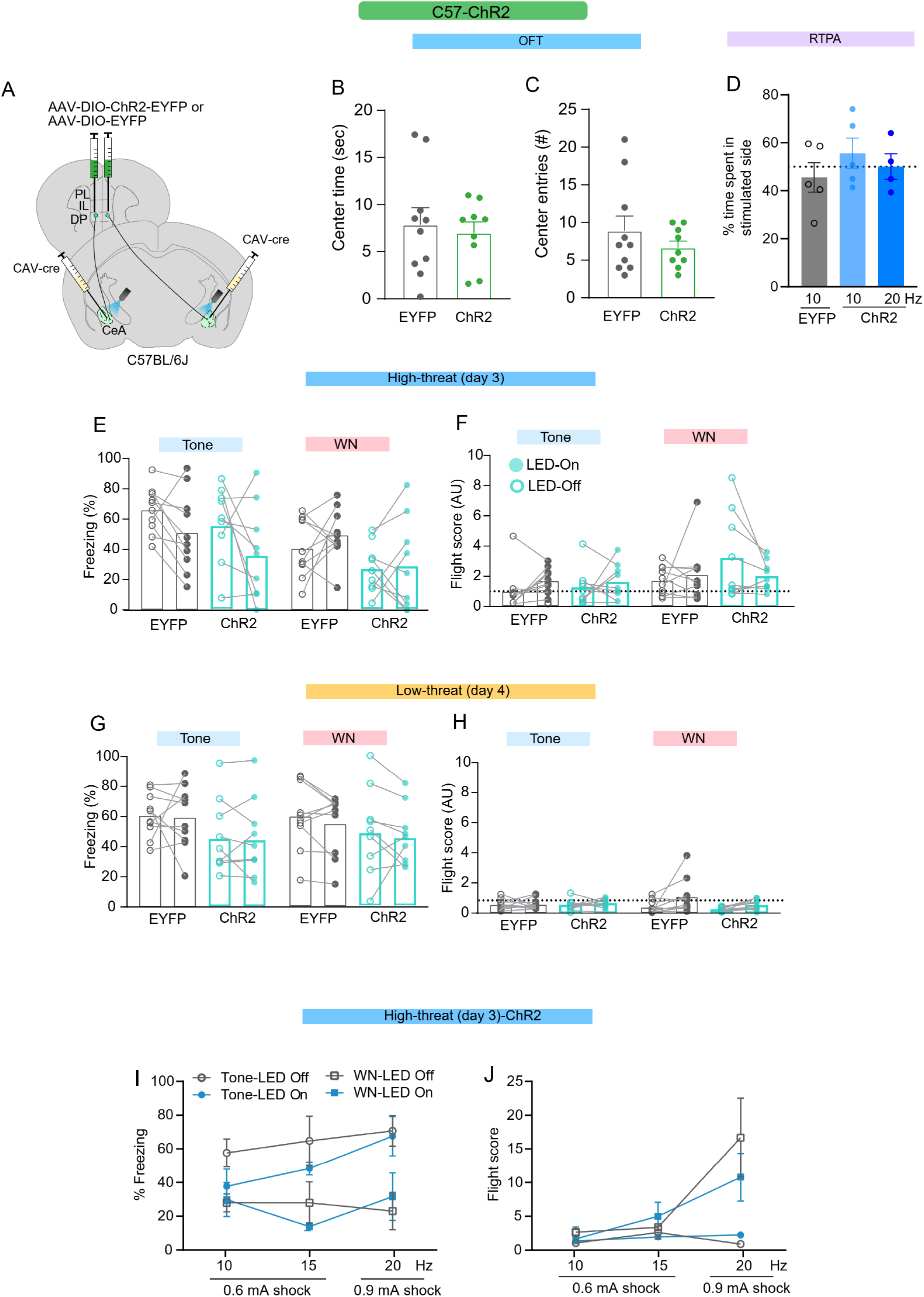
(Data related figure 4): Non-cell type specific stimulation of the DP-CEA pathway. **A,** Intersectional approach used to target optogenetic stimulation to DP-to-CeA terminals. **B-C**, Effect of optogenetic stimulation on OFT centre time (**B**) and centre entries (**C**) in EYFP (N = 10) and ChR2 (N = 9) groups (Unpaired t-test, n.s.). **D**, Effects of optogenetic stimulation on real-time place avoidance in EYFP (10 Hz, N = 5) and ChR2 (10 Hz, N = 5; 20 Hz, N = 4) groups (One-way ANOVA (F_(2, 11)_ = 0.73, *p* = 0.502). **E**, Effect of optogenetic excitation in EYFP (N = 10) and ChR2 (N = 9) groups on freezing in the high-threat context (LED-on vs LED-off, Mann-Whitney, n.s.). **F**, Effect of optogenetic excitation in EYFP (N = 10) and ChR2 (N = 9) groups on flight in the high-threat context (LED-on vs LED-off, Mann-Whitney, n.s.). **G**, Effect of optogenetic excitation in EYFP (N = 10) and ChR2 (N = 9) groups on freezing in the low-threat context (LED-on vs LED-off, Mann-Whitney, n.s.). **H**, Effect of optogenetic excitation in EYFP (N = 10) and ChR2 (N = 9) groups on flight in the low-threat context (LED-on vs LED-off, Mann-Whitney, n.s.). **I-J**, Freezing (**I**) and flight scores (**J**) during optogenetic stimulation during day 3 at different stimulation frequencies and shock intensities (at 0.6 mA - 10 Hz, N = 9; 15 Hz, N = 3; at 0.9 mA - 20 Hz, N = 5). Two-way ANOVA (*for % freezing*, Stimulation frequency x Shock intensity, F_(6, 56)_ = 0.76, *p* = 0.601, Stimulation frequency, F_(2, 56)_ = 1.10, *p* = 0.339, Shock intensity, F_(3, 56)_ = 8.37, *p* = 0.0001; *for flight*, Stimulation frequency x Shock intensity, F_(6, 56)_ = 4.42, *p* = 0.001, Shock intensity, F_(3, 56)_ = 6.66, *p* = 0.001, followed by Bonferroni’s post hoc test (tone/WN ON vs OFF non-significant).

**Figure S6.**
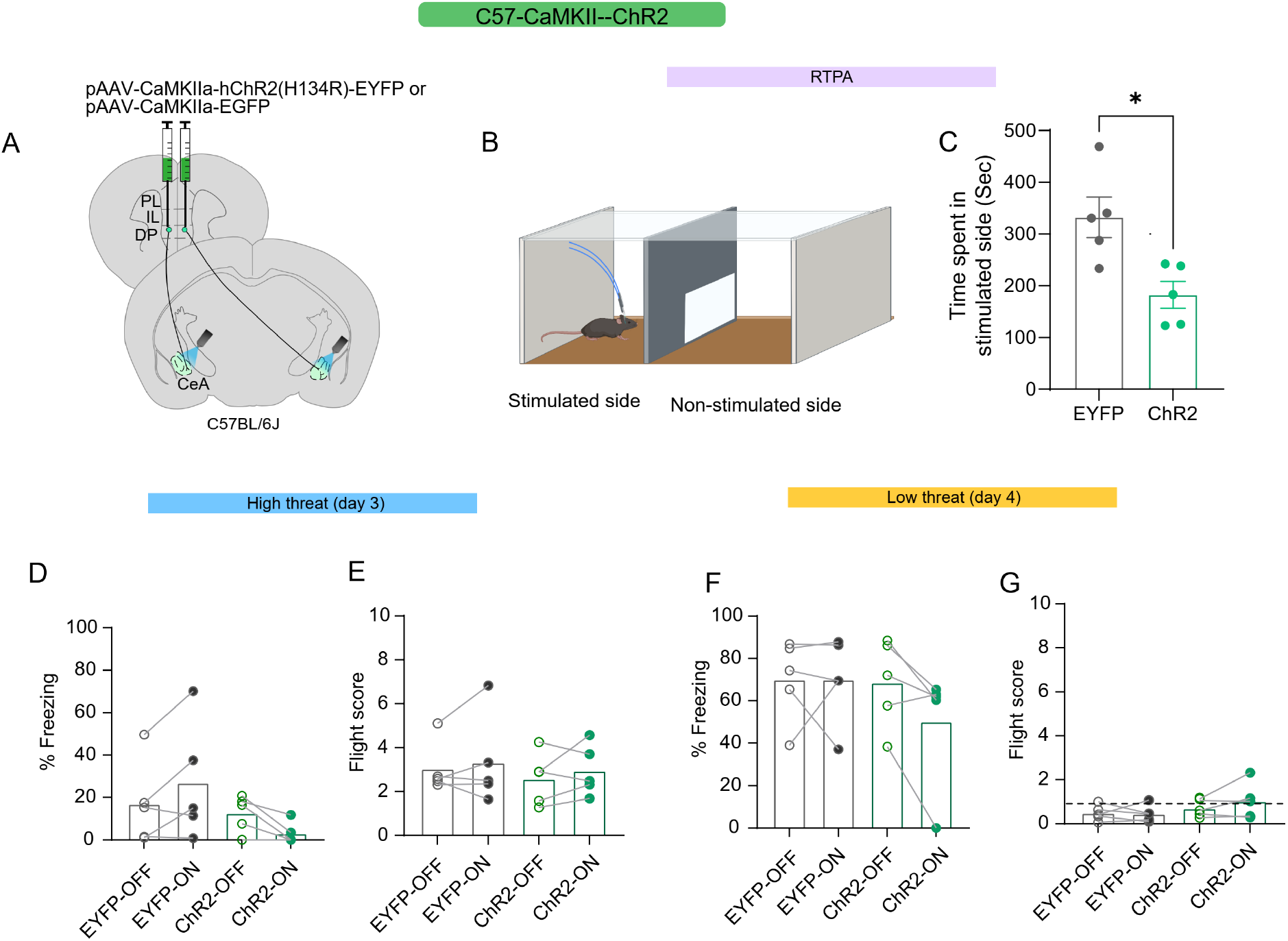
(Data related figure 4): Stimulation of theDP-CEA pathway using a CaMKII promotor. **A,** Viral injection strategy for optogenetic terminal stimulation of DP-to-CeA neuronal projections. **B-C**, Schematic (**B**) and results of real-time place aversion (RTPA) in EYFP (20 Hz, N = 5) and ChR2 (20 Hz, N = 5) groups. Unpaired t-test, t=3.191, df=8, *P <0.05. **D**, Effect of optogenetic excitation in EYFP (N = 5) and ChR2 (N = 5) groups on freezing in the high-threat context (LED-on vs LED-off, paired t-test, n.s.). **E**, Effect of optogenetic excitation in EYFP (N = 5) and ChR2 (N = 5) groups on flight scores in the high-threat context (LED-on vs LED-off, paired t-test, n.s.). **F**, Effect of optogenetic excitation in EYFP (N = 5) and ChR2 (N = 5) groups on freezing in the low-threat context (LED-on vs LED-off, paired t-test, n.s.). **G**, Effect of optogenetic excitation in EYFP (N = 5) and ChR2 (N = 5) groups on flight scores in the low-threat context (LED-on vs LED-off, paired t-test, n.s.).

**Figure S7.**
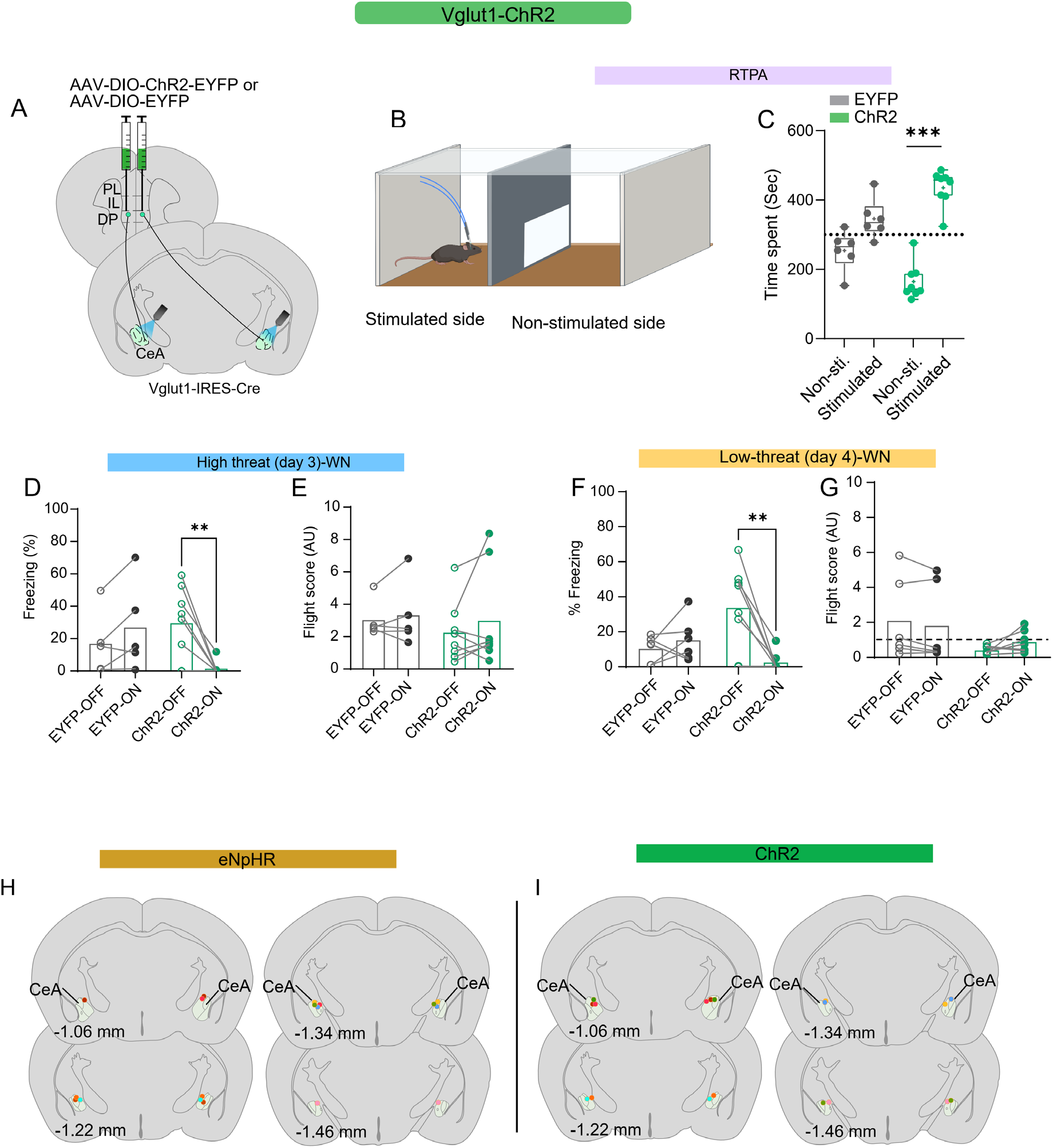
(Data related figure 4): Optogenetic stimulation of the DP-CEA pathway in Vglut1 mice. **A,** Viral injection strategy for optogenetic terminal stimulation of DP-to-CeA neuronal projections. **B-C**, Schematic (**B**) and real-time place aversion (RTPA) performance (**C**) from EYFP (N=6) and ChR2 (N=8) groups (unpaired t-test, EYFP (t=1.974, df=5, P =0.10), ChR2 (t=7.339, df=7, ***P =0.0002). **D**, Effect of optogenetic excitation in EYFP (N = 6) and ChR2 (N = 8) groups on freezing during WN in the high-threat context (LED-on vs LED-off, Paired t-test, ChR2, t=3.650, df=7, **P =0.008). **E**, Effect of optogenetic excitation in EYFP (N = 6) and ChR2 (N = 8) groups on flight during WN in the high-threat context (LED-on vs LED-off, Paired t-test, ChR2, t=1.077, df=7, P =0.31). **F**, Effect of optogenetic excitation in EYFP (N = 6) and ChR2 (N = 8) groups on freezing during WN in the low-threat context (LED-on vs LED-off, Paired t-test, ChR2, t=3.748, df=7, **P =0.007). **G**, Effect of optogenetic excitation in EYFP (N = 6) and ChR2 (N = 8) groups on flight score during WN in the low-threat context (LED-on vs LED-off, Paired t-test, ChR2, t=2.211, df=7, P =0.06). **H**, Example fibre placements over the CeA for the eNpHR groups (N = 9) **I**, Example fibre placements over the CeA for the ChR2 groups (N = 9)

**Figure S8.**
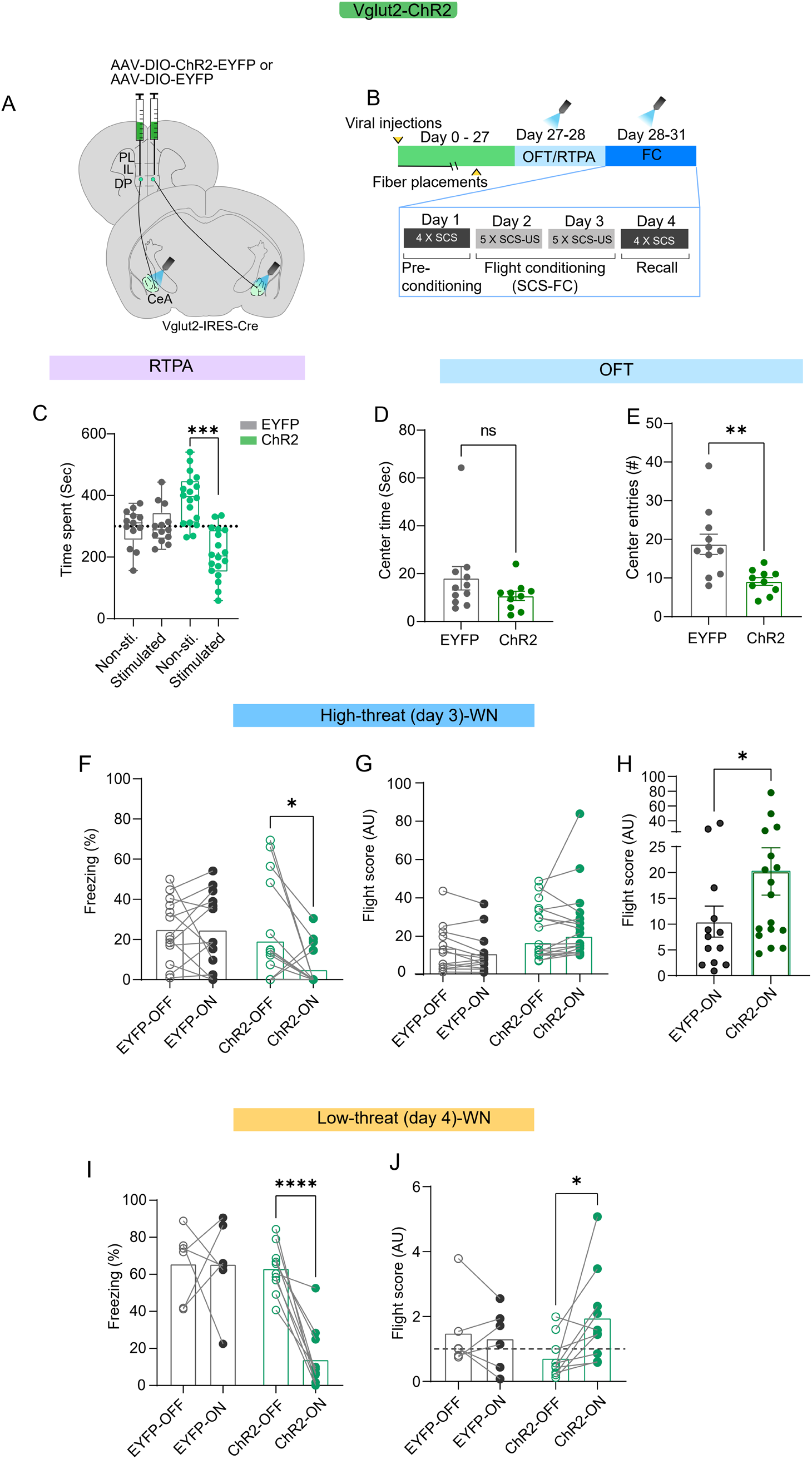
(Data related figure 4): Effect of DP-CEA pathway stimulation in Vglut2 mice. **A,** Viral injection strategy for optogenetic terminal stimulation of DP-to-CeA neuronal projections). **B**, Experimental timeline. **C**, Real-time place aversion (RTPA) performance of EYFP (N=13) and ChR2 (N=17) groups (paired t-test, EYFP (t=0.2167, df=12, P =0.83), ChR2 (t=4.713, df=17, ***P =0.0002). **D-E**, Effect of optogenetic excitation on OFT centre time (**D**) and number of entries into the centre zone (**E**) in EYFP (N=11) and ChR2 (N=10) groups (unpaired t-test (t=3.288, df=19, **P =0.01). **F**, Effect of optogenetic excitation in EYFP (N = 13) and ChR2 (N = 17) groups on freezing during WN in the high-threat context (LED-on vs LED-off, Paired t-test, ChR2, t=2.455, df=16, *P =0.025). **G**, Effect of optogenetic excitation in EYFP (N = 13) and ChR2 (N = 17) groups on flight during WN in the high-threat context (LED-on vs LED-off, paired t-test, ChR2, t=1.123, df=16). H, Comparison of flight scores in the LED-on condition between EYFP control and ChR2 groups (Mann Whitney test, \**P* = 0.0433). **I**, Effect of optogenetic excitation in EYFP (N = 6) and ChR2 (N = 10) groups on freezing during WN in the low-threat context (LED-on vs LED-off, paired t-test, ChR2, t=7.135, df=9, ****P <0.0001). **J**, Effect of optogenetic excitation in EYFP (N = 6) and ChR2 (N = 10) groups on flight scores during WN in the low-threat context (LED-on vs LED-off, paired t-test, ChR2, t=t=2.717, df=9, *P = 0.02).

**Figure S9:**
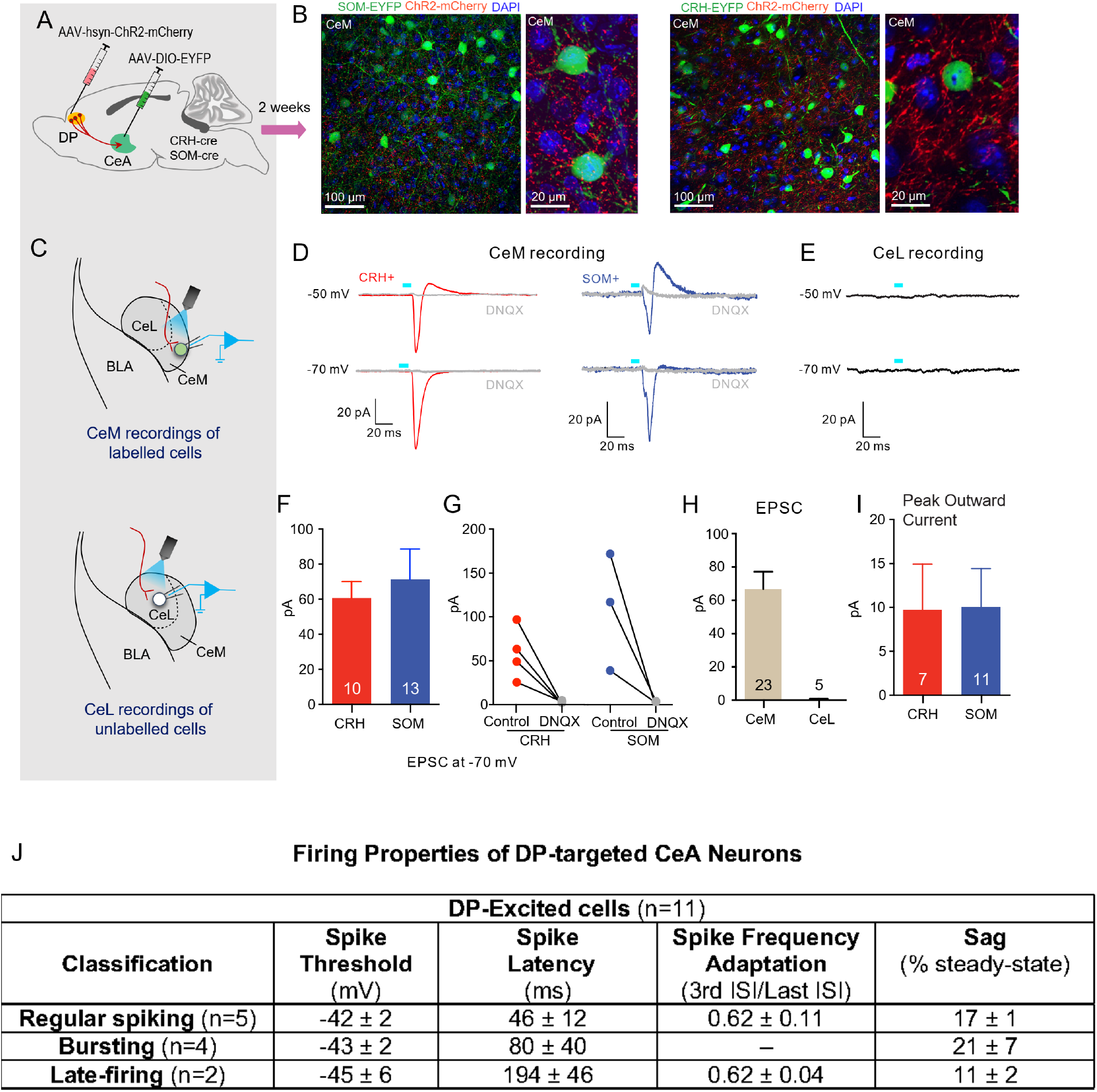
Data related to Figure 5. **A**, Schematic of targeting strategy for optogenetic stimulation. **B**, DP terminals in CeM near SOM+ (*left*) and CRH+ (*right*) cells at 20x and 40x magnification. **C**, Strategy for recording light-evoked synaptic input from DP to SOM+ or CRH+ neurons from CeM (*top*) and CeL (*bottom*) regions. **D**, Representative evoked synaptic responses in CeM SOM+ and CRH+ cells by photostimulation of DP axonal fibres in voltage-clamp. **E**, Photostimulation of axonal fibres did not evoke responses in CeL neurons. **F**, Average amplitude of evoked EPSCs in CRH+ neurons (N = 10 cells from 3 mice) and SOM+ (N = 13 cells from 3 mice) at -70mV. Unpaired Student’s t-test, *p* = 0.63. **G**, Amplitudes of evoked EPSCs in CRH+ (N = 4 cells from 3 mice) and SOM+ (N = 3 cells from 2 mice) neurons at -70 mV, before and after application of DNQX. **H**, Average amplitude of evoked EPSCs in CeM (N = 23 cells from 6 mice) and CeL neurons (N = 5 cells from 2 mice). **I**, The amplitude of disynaptic IPSCs evoked by ChR2 stimulation of DP terminals (Unpaired Student’s t-test, t_(16)_= 0.055; p = 0.96). **J,** The firing properties of DP-targeted CeM neurons.

## Notes

### Competing Interest Statement

The authors have declared no competing interest.

### Summary of Updates

Extensive revision of all figures and main text. Author affiliations updated and supplemental figures updated.

## References

1. Fonzo, G. A. et al. PTSD psychotherapy outcome predicted by brain activation during emotional reactivity and regulation. Am. J. Psychiatry 174, 1163–1174 (2017).

2. Fadok, J. P. et al. A competitive inhibitory circuit for selection of active and passive fear responses. Nature 542, 96–99 (2017).

3. Roelofs, K. Freeze for action: Neurobiological mechanisms in animal and human freezing. *Philos. Trans. R. Soc. Lond., B*, Biol. Sci. 372 (2017).

4. Fanselow, M. S., Hoffman, A. N. & Zhuravka, I. Timing and the transition between modes in the defensive behavior system. Behav. Processes 166, 103890 (2019).

5. Blanchard, D. C. & Blanchard, R. J. Defensive behaviors, fear, and anxiety. In Handbook of anxiety and fear. 63–79 (Elsevier Academic Press, 2008). doi:10.1016/S1569-7339(07)00005-7.

6. Perusini, J. N. & Fanselow, M. S. Neurobehavioral perspectives on the distinction between fear and anxiety. Learn. Mem. 22, 417–425 (2015).

7. Johnson, P. L., Truitt, W. A., Fitz, S. D., Lowry, C. A. & Shekhar, A. Neural pathways underlying lactate-induced panic. Neuropsychopharmacol. 33, 2093–2107 (2008).

8. Münsterkötter, A. L. et al. Spider or no spider? Neural correlates of sustained and phasic fear in spider phobia. Depress. Anxiety 32, 656–663 (2015).

9. Mobbs, D. et al. From threat to fear: The neural organization of defensive fear systems in humans. J. Neurosci. 29, 12236–12243 (2009).

10. Tromp, D. P. M. et al. Reduced structural connectivity of a major frontolimbic pathway in generalized anxiety disorder. Arch. Gen. Psychiatry 69, 925–934 (2012).

11. Marek, R., Strobel, C., Bredy, T. W. & Sah, P. The amygdala and medial prefrontal cortex: Partners in the fear circuit. J. Physiol. 591, 2381–2391 (2013).

12. Senn, V. et al. Long-range connectivity defines behavioral specificity of amygdala neurons. Neuron 81, 428–437 (2014).

13. Karalis, N. et al. 4-Hz oscillations synchronize prefrontal-amygdala circuits during fear behavior. Nat. Neurosci. 19, 605–612 (2016).

14. Andrewes, D. G. & Jenkins, L. M. The role of the amygdala and the ventromedial prefrontal cortex in emotional regulation: Implications for post-traumatic stress disorder. Neuropsychol. Rev. 29, 220–243 (2019).

15. De Franceschi, G., Vivattanasarn, T., Saleem, A. B. & Solomon, S. G. Vision guides selection of freeze or flight defense strategies in mice. Curr. Biol. 26, 2150–2154 (2016).

16. Wang, W. et al. Coordination of escape and spatial navigation circuits orchestrates versatile flight from threats. Neuron 109, 1–13 (2021).

17. Mcdonald, A. J. Cortical pathways to the mammalian amygdala. Prog. Neurobiol. 55, 257–332 (1998).

18. Kataoka, N., Shima, Y., Nakajima, K. & Nakamura, K. A central master driver of psychosocial stress responses in the rat. Science 367, 1105–1112 (2020).

19. Anastasiades, P. G. & Carter, A. G. Circuit organization of the rodent medial prefrontal cortex. Trends Neurosci. 44, 550–563 (2021).

20. Fremeau, R. T. et al. The expression of vesicular glutamate transporters defines two classes of excitatory synapse. Neuron 31, 247–260 (2001).

21. Borkar, C. D. & Fadok, J. P. A novel pavlovian fear conditioning paradigm to study freezing and flight behavior. J Vis Exp 1–13 (2021). doi:10.3791/61536.

22. Anderson, D. J. & Adolphs, R. A framework for studying emotions across species. Cell 157, 187–200 (2014).

23. Fadok, J. P., Markovic, M., Tovote, P. & Lüthi, A. New perspectives on central amygdala function. Curr. Opin. Neurobiol. 49, 141–147 (2018).

24. Dumont, É. C., Martina, M., Samson, R. D., Drolet, G. & Paré, D. Physiological properties of central amygdala neurons: Species differences. Eur. J. Neurosci. 15, 545–552 (2002).

25. Duvarci, S., Popa, D. & Paré, D. Central amygdala activity during fear conditioning. J. Neurosci. 31, 289–294 (2011).

26. Li, J. N. & Sheets, P. L. The central amygdala to periaqueductal gray pathway comprises intrinsically distinct neurons differentially affected in a model of inflammatory pain. J. Physiol. 596, 6289–6305 (2018).

27. Rizvi, T. A., Ennis, M., Behbehani, M. M. & Shipley, M. T. Connections between the central nucleus of the amygdala and the midbrain periaqueductal gray: Topography and reciprocity. J. Comp. Neurol. 303, 121–131 (1991).

28. Tovote, P. et al. Midbrain circuits for defensive behaviour. Nature 534, 206–212 (2016).

29. Bandler, R. & Carrive, P. Integrated defence reaction elicited by excitatory amino acid microinjection in the midbrain periaqueductal grey region of the unrestrained cat. Brain Res. 439, 95–106 (1988).

30. Behbehani, M. M. Functional characteristics of the midbrain periaqueductal gray. Prog. Neurobiol. 46, 575–605 (1995).

31. Keifer, O. P., Hurt, R. C., Ressler, K. J. & Marvar, P. J. The physiology of fear: Reconceptualizing the role of the central amygdala in fear learning. Physiology 30, 389– 401 (2015).

32. Ressler, R. L. & Maren, S. Synaptic encoding of fear memories in the amygdala. Curr. Opin. Neurobiol. 54, 54–59 (2019).

33. Kong, M. S. & Zweifel, L. S. Central amygdala circuits in valence and salience processing. Behav. Brain Res. 410, 113355 (2021).

34. Viviani, D. et al. Oxytocin selectively gates fear responses through distinct outputs from the central amygdala. Science 333, 104–107 (2011).

35. Evans, DA., et al. A synaptic threshold mechanism for computing escape decisions. Nature 558, 46–76 (2009).

36. Wang, W. et al. Dorsal premammillary projection to periaqueductal gray controls escape vigor from innate and conditioned threats. Elife 10, 1–30 (2021).

37. Li, H. et al. Experience-dependent modification of a central amygdala fear circuit. Nat. Neurosci. 16, 332–339 (2013).

38. Hunt, S., Sun, Y., Kucukdereli, H., Klein, R. & Sah, P. Intrinsic circuits in the CeL. eNeuro 4, 1–18 (2017).

39. Tovote, P., Fadok, J. P. & Lüthi, A. Neuronal circuits for fear and anxiety. Nat. Rev. Neurosci. 16, 317–331 (2015).

40. Quirk, G. J., Likhtik, E., Pelletier, J. G. & Paré, D. Stimulation of medial prefrontal cortex decreases the responsiveness of central amygdala output neurons. J. Neurosci. 23, 8800–8807 (2003).

41. Bukalo, O. et al. Prefrontal inputs to the amygdala instruct fear extinction memory formation. Sci. Adv. 1, 1–9 (2015).

42. Hersman, S., Allen, D., Hashimoto, M., Brito, S. I. & Anthony, T. E. Stimulus salience determines defensive behaviors elicited by aversively conditioned serial compound auditory stimuli. Elife 9, (2020).

43. Dong, P. et al. A novel cortico-intrathalamic circuit for flight behavior. Nat. Neurosci. 22, 941–949 (2019).

44. Totty, M. S. et al. Behavioral and brain mechanisms mediating conditioned flight behavior in rats. Sci. Rep. 11, 1–15 (2021).

## Methods References

45. Soudais, C., Laplace-Builhe, C., Kissa, K. & Kremer, E. J. Preferential transduction of neurons by canine adenovirus vectors and their efficient retrograde transport in vivo. FASEB J. 15, 2283–2285 (2001).

46. Resendez, S. L. et al. Visualization of cortical, subcortical and deep brain neural circuit dynamics during naturalistic mammalian behavior with head-mounted microscopes and chronically implanted lenses. Nat. Protoc. 11, 566–597 (2016).

47. Corder, G. et al. An amygdalar neural ensemble that encodes the unpleasantness of pain. Science 363, 276–281 (2019).

48. Ghosh, K. K. et al. Miniaturized integration of a fluorescence microscope. Nat. Methods 8, 871–878 (2011).

49. Parker, J. G. et al. Diametric neural ensemble dynamics in parkinsonian and dyskinetic states. Nature 557, 177–182 (2018).

50. Chen, C. et al. Astrocytes amplify neuronal dendritic volume transmission stimulated by norepinephrine. Cell Rep. 29, 4349–4361.e4 (2019).

